# Mitochondrial copper stabilizes lipoylated TCA cycle proteins to sustain metabolism and proliferation

**DOI:** 10.64898/2026.05.20.726656

**Authors:** Santanu Ghosh, Andrew F. Jarvis, Jordi C.J. Hintzen, Nate R. McKnight, Pedro Costa-Pinheiro, Noelle H. Noughton, Yumi Kim, Alison Jaccard, Scot C. Leary, Paul A. Cobine, Caroline R. Bartman, Gina M. DeNicola, Nathaniel W. Snyder, Kathryn E. Wellen, George M. Burslem, Donita C. Brady

## Abstract

Copper (Cu) is an essential cofactor for mitochondrial cytochrome *c* oxidase, yet whether it directly regulates mitochondrial metabolism beyond respiration remains unclear. Here we show that mitochondrial Cu, delivered by SLC25A3, is required to maintain the stability of lipoylated TCA cycle proteins. Loss of *Slc25a3* or pharmacological Cu depletion selectively destabilized the lipoylated E2 subunits of mitochondrial dehydrogenases and the lipoylation enzymes LIPT1 and LIPT2, an effect not reproduced by acute electron transport chain inhibition. Mechanistically, we find that Cu directly engages the reduced lipoyl moiety using chemical probes and synthetic peptide approaches. Cu depletion impaired PDH and OGDH activity, rewired TCA cycle metabolism, and imposed a dependence on pyruvate carboxylase for anaplerosis. This metabolic defect depleted aspartate, suppressed mTORC1 signaling, and limited proliferation. Conversely, selective delivery of Cu to the mitochondria restored lipoylation, TCA cycle function, and cell growth. Together, these findings identify mitochondrial Cu as a structural regulator of the lipoylation machinery and reveal a direct link between Cu homeostasis and central carbon metabolism.

## Introduction

Copper (Cu) is an essential transition metal whose redox properties support the activity of enzymes such as cytochrome *c* oxidase (COX) and superoxide dismutase^1,2^. Disruption of Cu homeostasis causes severe disease, including neurodevelopmental defects in Menkes disease and hepatic Cu accumulation with progressive liver and neurological dysfunction in Wilson disease, underscoring its importance in cellular physiology^3,4^. Beyond these canonical roles, emerging evidence indicates that Cu can act as a dynamic regulator of signaling and metabolism through direct interactions with protein targets, expanding its function beyond that of a static catalytic cofactor^5,6^.

Cu homeostasis is tightly controlled at the subcellular level, with dedicated transporters and chaperones serving to establish and maintain distinct Cu pools in the Golgi apparatus, cytosol, and mitochondria^7,8^. This compartmentalization enables Cu to be selectively delivered to specific protein targets while limiting toxicity. Within mitochondria, Cu is primarily required for the assembly and activity of COX [Complex IV of the electron transport chain (ETC)], and a labile matrix Cu pool has been described^9–11^. The dual Cu and phosphate carrier SLC25A3 was identified as a mitochondrial Cu transporter required for COX biogenesis, providing a direct route for Cu entry into the mitochondrial matrix^12^. Mutations in *SLC25A3* in humans cause mitochondrial disease characterized by defects in oxidative phosphorylation, cardiomyopathy, and neuromuscular dysfunction, highlighting the importance of mitochondrial Cu homeostasis^13,14^. However, whether mitochondrial Cu regulates metabolic processes beyond its role in respiration remains unclear.

Mitochondrial respiration is tightly coupled to proliferative metabolism, not simply through ATP production, but through its role in sustaining biosynthetic pathways. Seminal studies demonstrated that a primary function of the ETC in proliferating cells is to provide electron acceptors required for aspartate synthesis, redefining the role of respiration in cell growth^15,16^. Consistent with this, ETC inhibition leads to aspartate depletion, activation of the integrated stress response, and suppression of mTORC1 signaling, thereby limiting anabolic growth and proliferation^17,18^. Recent work further suggests that perturbations in Cu availability can impact mitochondrial metabolism and nucleotide synthesis^19^. Together, these findings establish mitochondrial function as a central regulator of metabolic and growth signaling networks.

Recent work has further linked Cu to mitochondrial metabolism through the discovery of cuproptosis, a Cu-dependent cell death pathway induced by Cu ionophores such as elesclomol^20^. In this context, pharmacologically induced mitochondrial Cu accumulation directly engages lipoylated components of the TCA cycle, promoting their aggregation and proteotoxic stress. These observations identify lipoylated mitochondrial proteins as Cu-responsive targets, but whether physiological Cu levels within the organelle regulate their stability and function under non-lethal conditions has yet to be clarified.

Here, we show that mitochondrial Cu, specifically Cu(I), delivered by SLC25A3, is required to stabilize lipoylated TCA cycle proteins and sustain mitochondrial metabolism. Loss of mitochondrial Cu selectively destabilizes lipoylated dehydrogenase complexes, impairing oxidative TCA cycle metabolism and imposing a dependence on pyruvate carboxylase (PC)-mediated anaplerosis. This metabolic rewiring depletes aspartate, suppresses mTORC1 signaling, and limits proliferation. Conversely, restoration of mitochondrial Cu rescues lipoylation, metabolic function, and cell growth. Together, these findings identify mitochondrial Cu as a direct regulator of the lipoylation machinery and establish a mechanistic link between Cu homeostasis, mitochondrial metabolism, and proliferative signaling.

## Results

### *Slc25a3* loss impairs mitochondrial respiration and reduces mTORC1 signaling

To determine whether mitochondrial Cu directly supports respiratory function independently of phosphate transport, we disrupted *Slc25a3* in mouse embryonic fibroblasts (MEFs) and reconstituted cells with either wild-type SLC25A3 (WT) or a Cu-transport-competent, phosphate-deficient mutant (L175A; MUT)^21^ (Extended Data Fig. 1a). Loss of *Slc25a3* abolished COX activity, which was restored by both WT and mutant SLC25A3 (Extended Data Fig. 1b), indicating that mitochondrial Cu delivery is required for COX function^12,22^.

To determine whether mitochondrial Cu supports respiratory function linked to aspartate-dependent growth signaling (Fig. 1a), we assessed ETC activity following loss of *Slc25a3*. Oxygen consumption rate (OCR) analysis revealed reduced responses to oligomycin, FCCP, and rotenone/antimycin A (Fig. 1b), indicating impaired ETC function^23^. WT SLC25A3 restored respiration, whereas the mutant partially rescued this defect (Fig. 1b). In parallel, mitochondrial NAD⁺/NADH ratios were reduced and restored upon mutant reconstitution (Fig. 1c), indicating that Cu transport is sufficient to support mitochondrial redox balance^24^.

**Figure 1.**
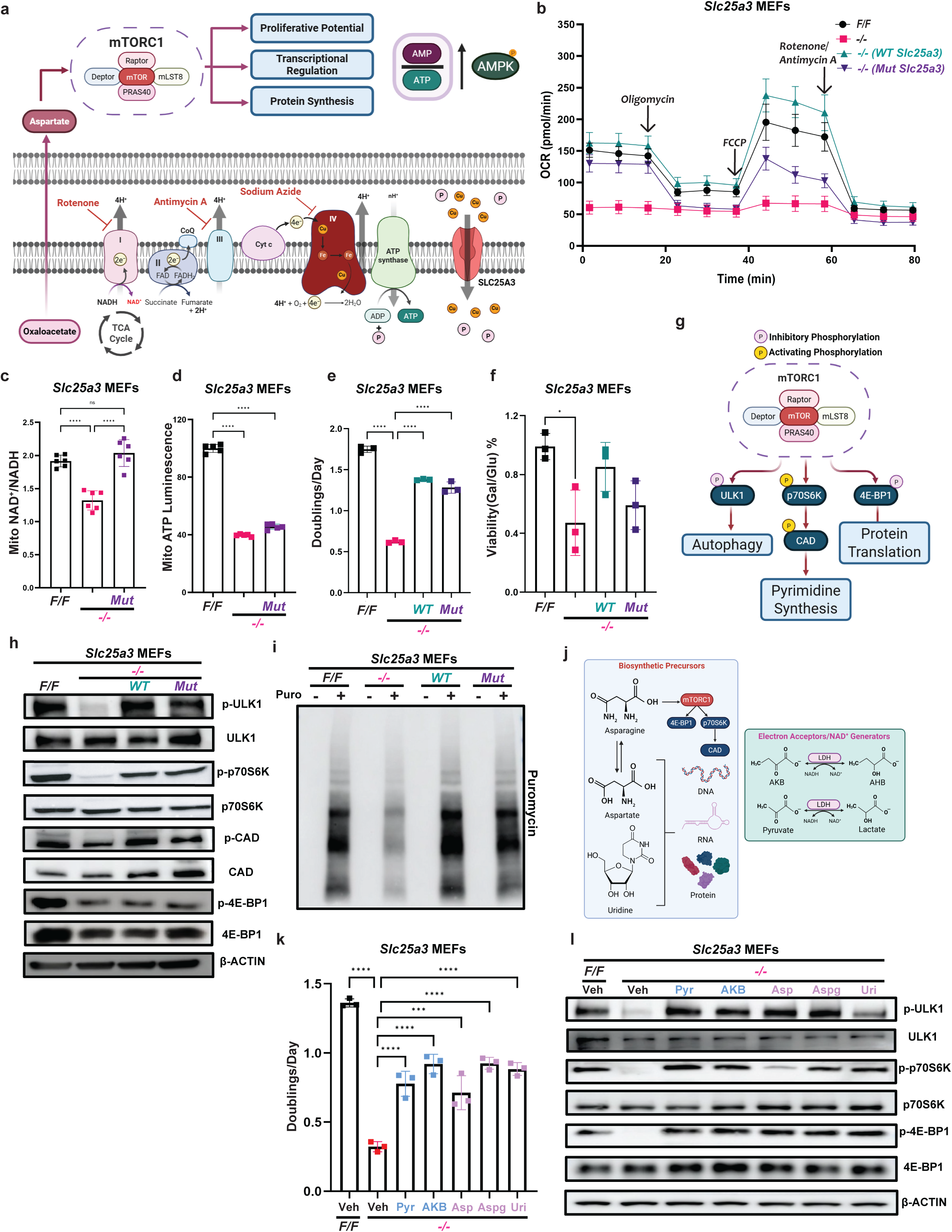
Slc25a3 loss impairs mitochondrial respiration and reduces mTORC1 signaling. **a,** Schematic representation of the effects of copper dysregulation in the mitochondria. Copper enters the mitochondrial matrix through the *Slc25a3* transporter and is necessary for the proper functioning of the ETC and TCA cycle. Oxaloacetate produced by the TCA is converted into aspartate which positively regulates the mTORC1 complex for various biologically important functions. The *Slc25a3* transporter also transports phosphate into the matrix for oxidative phosphorylation and the generation of ATP. Disruption of ATP production leads to the activation of AMPK and subsequent catabolic processes. **b,** Oxygen consumption rate measurements over time with sequential injections of 1.5µM oligomycin, 1µM FCCP and 0.5µM Rotenone/Antimycin A, in *Slc25a3 F/F*, *-/-*, *WT* and *Mut* MEFs. **c,** NAD^+^/NADH ratios from isolated mitochondria extracted from *Slc25a3 F/F*, *-/-* and *Mut* MEFs. **d,** Relative ATP levels measured in isolated mitochondria from *Slc25a3 F/F*, *-/-* and *Mut* MEFs. **e,** Doublings per day calculated from cell counts measured after 48 hours using trypan blue dye exclusion in *Slc25a3 F/F*, *-/-*, *WT* and *Mut* MEFs. **f,** Viability (% of cells alive) of *Slc25a3 F/F*, *-/-*, *WT* and *Mut* MEFs in 10mM Galactose and 10mM Glucose, for 24 hours, measured using trypan blue dye exclusion, represented as a ratio of viability in galactose over viability in glucose. **g,** Schematic representation of the mTORC1 signaling pathway and its downstream phenotypic effects **h,** Immunoblot detection of p-ULK1 (Ser757), total ULK1, p-p70S6K (Thr389), total p70S6K, p-CAD (Ser1859), total CAD, p-4E-BP1 (Ser65), total 4E-BP1, or β-ACTIN in *Slc25a3 F/F*, *-/-*, *WT* and *Mut* MEFs. **i,** Immunoblot detection of puromycin in *Slc25a3 F/F*, *-/-*, *WT* and *Mut* MEFs treated with or without 10mg/mL puromycin for 15 minutes. **j,** Diagrammatic representation of biosynthetic precursors and NAD^+^ generators used to rescue the effects of ETC inhibition **k, l, (k)** Doublings per day calculated from cell counts measured using trypan blue dye exclusion or **(m)** immunoblot detection of p-ULK1, total ULK1, p-p70S6K, total p70S6K, p-4E-BP1, total 4E-BP1, or β-ACTIN in *Slc25a3 F/F* MEFs treated with vehicle, or *Slc25a3 -/-* MEFs treated with either vehicle, 20mM Aspartate (Asp), 20mM Asparagine (Aspg), 1mM Uridine (Uri), 2mM Pyruvate (Pyr) or 1mM α-ketobutyrate (AKB). For **c-f**, **h-i**, **k** and **l** n= 3 biologically independent experiments. For **b-f** and **k**, data represented as mean ± SEM. ns= not significant, *p≤ 0.05, ***p≤ 0.001, ****p≤ 0.0001.

This defect in respiration was accompanied by altered cellular energetics. Total ATP levels were reduced upon loss of *Slc25a3* and restored by mutant SLC25A3 (Extended Data Fig. 1c), whereas mitochondrial ATP production remained reduced in mutant-rescued cells (Fig. 1d). Additionally, separation of the glycolytic and mitochondrial ATP production rates revealed that oxditative phosphorylation is highly compromised in cells lacking the SLC25A3 transporter whereas glycolytic rates remained slightly elevated, likely as a compensatory feature (Extended Data Fig. 1d). Consistent with this imbalance, AMPK phosphorylation was increased (Extended Data Fig. 1e), and the abundance of ETC complex subunits was reduced (Extended Data Fig. 1f), indicating disruption of mitochondrial function^25,26^.

Given these changes in mitochondrial activity, we examined how cells adapt metabolically. Loss of *Slc25a3* was associated with increased expression of glycolytic genes (Extended Data Fig. 1g), along with elevated glucose consumption (Extended Data Fig. 1h) and lactate secretion (Extended Data Fig. 1i), consistent with a shift toward glycolysis^27,28^. However, this adaptation was insufficient to sustain proliferation, as cells lacking *Slc25a3* exhibited reduced growth rates (Fig. 1e) and impaired viability under galactose conditions that require oxidative metabolism^29^ (Fig. 1f). Re-expression of the phosphate-deficient mutant of SLC25A3 only partially rescued viability in galactose owing to ETC inhibition under phosphate-depleted conditions^30^. These defects were not observed in MEFs lacking the high-affinity Cu transporter *Slc31a1*, which mediates the majority of cellular Cu uptake^31,32^ (Extended Data Fig. 1j,k), indicating that they arise specifically from loss of mitochondrial Cu.

Because mitochondrial respiration supports aspartate-dependent growth signaling, we next examined mTORC1 activity (Fig. 1g). Loss of *Slc25a3* decreased phosphorylation of ULK1, p70S6K, CAD, which is downstream of p70S6K, and 4E-BP1^17,33,34^ (Fig. 1g,h). These changes were restored by SLC25A3 re-expression but not observed in *Slc31a1*⁻*^/^*⁻ MEFs (Extended Data Fig. 1l), indicating that mTORC1 suppression results from mitochondrial Cu depletion. Downstream of mTORC1 signaling, we also observed reduced protein synthesis which re-expression of SLC25A3 similarly restored (Fig. 1i), consistent with impaired anabolic signaling^35,36^.

To determine whether these effects reflect a general response to ETC inhibition, we compared cells lacking *Slc25a3* with cells treated with inhibitors of Complex I, III, or IV. RNA-seq analysis revealed that loss of *Slc25a3* induces a transcriptional state distinct from all acute ETC inhibition conditions (Extended Data Fig. 2a), with upregulation of hypoxia-related pathways and downregulation of oxidative phosphorylation and mTORC1 signaling gene sets^37,38^ (Extended Data Fig. 2b-h).

Finally, we tested whether restoring key metabolic intermediates could rescue proliferation (Fig. 1j). Supplementation with aspartate (Asp), asparagine (Aspg), uridine (Uri), pyruvate (Pyr), or α-ketobutyrate (AKB) partially restored growth (Fig. 1k) and mTORC1 signaling (Fig. 1l), but did not fully rescue proliferation, indicating that loss of mitochondrial Cu imposes both mTORC1-dependent and independent constraints on cell growth^16,17,39^. Together, these data demonstrate that loss of mitochondrial Cu disrupts ETC function, induces compensatory metabolic rewiring, and suppresses mTORC1-dependent growth signaling, thereby impairing cell proliferation.

### Mitochondrial Cu loss rewires TCA cycle metabolism through impaired dehydrogenase activity

Given the respiratory defect observed upon loss of *Slc25a3*, we first asked whether impaired ETC function alone could account for the proliferative phenotype. To test this, we restored electron transport or redox balance using alternative oxidase (AOX), NDI1, and cytosolic or mitochondrial NADH oxidases (LbNOX and mitoLbNOX)^40–42^ (Extended Data Fig. 3a,b). Although these interventions partially improved proliferation, none restored growth to control levels (Extended Data Fig. 3c). Similarly, pharmacological approaches targeting redox balance like nicotinic acid (NA) and nicotinamide (NAM) which are precursors for NAD^+^ synthesis, lactate metabolism like the Lactate Dehydrogenase (LDH) inhibitor (GSK2837808A), TCA cycle intermediates like sodium propionate (Na-propionate) which can enter the TCA cycle through succinyl-CoA, or reactive oxygen species like N-acetylcysteine (NAC) which helps produce more GSH, failed to rescue proliferation (Extended Data Fig. 3d-g). These results indicate that the growth defect associated with loss of *Slc25a3* cannot be explained solely by ETC dysfunction or redox imbalance.

We therefore examined whether mitochondrial Cu loss alters metabolite abundance. Unbiased metabolomic profiling revealed a distinct metabolic signature in cells lacking *Slc25a3* that differed from that observed following acute inhibition of Complex I, III, or IV (Extended Data Fig. 4a,b), indicating that the metabolic consequences of *Slc25a3* loss are not simply due to ETC blockade^43,44^. Specifically, cells lacking *Slc25a3* exhibited accumulation of pyruvate and α-ketoglutarate (α-KG), together with depletion of acetyl-CoA, succinate, fumarate, malate, oxaloacetate, and aspartate (Fig. 2a). Notably, these cells also showed increased levels of 2-hydroxyglutarate (Extended Data Fig. 2c,d), suggesting that reductive stress arising from disrupted mitochondrial NAD+/NADH homeostasis (Fig. 1c) drives the aberrant reduction of α-ketoglutarate to 2-hydroxyglutarate^45^. In contrast, acute inhibition of Complex IV resulted in accumulation of TCA intermediates (Fig. 2b), further supporting that loss of Slc25a3 induces metabolic changes beyond ETC dysfunction. Further, global analysis of differentially abundant metabolites revealed coordinated alterations across nucleotide, amino acid, and central carbon metabolism pathways (Extended Data Fig. 4a), indicating broad metabolic reprogramming^46^.

**Figure 2.**
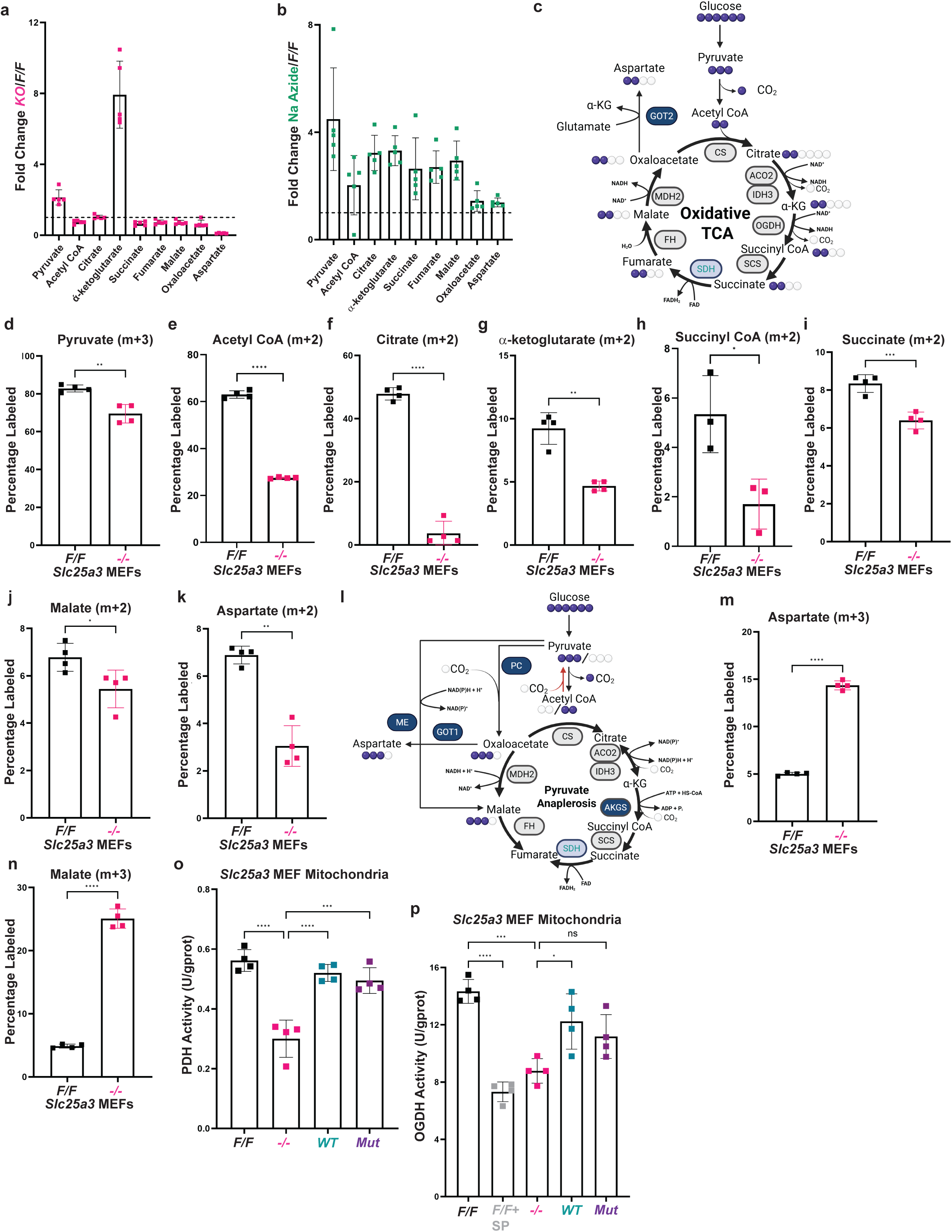
Mitochondrial copper loss rewires TCA cycle metabolism through impaired dehydrogenase activity. **a, b,** Relative abundance of steady-state untargeted metabolites in **(a)** *Slc25a3 -/-* MEFs relative to *Slc25a3 F/F* MEFs or **(b)** *Slc25a3 F/F* cells treated with 500nM Sodium Azide relative to treatment with vehicle, represented as fold change. The dashed line represents a fold change of 1, indicating no change in the metabolite levels in the treatment conditions. **c,** Schematic of the ^13^C-glucose tracing pathway starting from glucose to Acetyl-CoA, which enters the TCA and cycles in the forward clockwise/oxidative direction, consuming NAD^+^, producing oxaloacetate which is converted to aspartate using glutamic-oxaloacetic transaminase 2 (GOT2). The purple spheres represent ^13^C-labeled carbon atoms while the white spheres represent ^12^C-labeled carbon atoms. **d-k,** ^13^C-glucose (10mM) stable isotope tracing (6 hour pulse) labeling into **(d)** m+3 pyruvate, **(e)** m+2 acetyl CoA, **(f)** m+2 citrate, **(g)** m+2 α-ketoglutarate, **(h)** m+2 succinyl CoA, **(i)** m+2 succinate, **(j)** m+2 malate, and **(k)** m+2 aspartate, in extracts from *Slc25a3 F/F* and *-/-* MEFs, as measured by GC-MS or LC-MS(acetyl- and succinyl-CoA). **l,** Schematic of ^13^C-glucose tracing pathway from glucose to pyruvate, which enters the TCA cycle through pyruvate carboxylase (PC) and proceeds in an anti-clockwise/reductive direction, typically seen in hypoxic or ETC-inhibited cells. Pyruvate can also enter the TCA through malic enzyme which converts pyruvate directly to malate. This pathway generates m+3 states of the TCA cycle metabolites, as well as aspartate, as opposed to m+2 states generated by the oxidative TCA. **m, n,** ^13^C-glucose (10mM) stable isotope tracing (6 hour pulse) labeling into **(m)** m+3 aspartate, and **(n)** m+3 malate, in extracts from *Slc25a3 F/F* and *-/-* MEFs, as measured by GC-MS. **o,** Pyruvate Dehydrogenase activity measured using an activity assay kit in isolated mitochondria from *Slc25a3 F/F*, *-/-*, *WT* and *Mut* MEFs, normalized to respective protein concentrations. **p,** 2-Oxoglutarate Dehydrogenase activity measured using an activity assay kit in isolated mitochondria from *Slc25a3 F/F* MEFs treated with either vehicle or succinyl phosphonate (100µM), *Slc25a3 -/-*, *WT* and *Mut* MEFs, normalized to respective protein concentrations. For **o** and **p**, n=3 biologically independent experiments. For **a**, **b**, **d-k**, **m-p** data represented as mean ± SEM. ns= not significant, *p≤ 0.05, **p≤0.01, ***p≤ 0.001, ****p≤ 0.0001.

To define the underlying metabolic defect, we traced carbon labeling through the TCA cycle using uniformly labeled ¹³C-glucose^47,48^ (Fig. 2c). Incorporation of glucose-derived carbon into pyruvate (m+3 isotopologue) was somewhat reduced (Fig. 2d), and acetyl-CoA, citrate, α-KG, succinyl-CoA, and succinate (m+2 isotopologues) was markedly reduced upon loss of *Slc25a3* (Fig. 2e-i), indicating impaired entry of pyruvate into the oxidative TCA cycle. Labeling of downstream intermediates and aspartate was also reduced (Fig. 2j,k), consistent with decreased oxidative TCA cycle metabolism.

In contrast, analysis of m+3 isotopologues revealed increased labeling of malate, and aspartate (Fig. 2l-n), indicating engagement of an alternative anaplerotic route into the TCA cycle. The m+3 labeling pattern is consistent with pyruvate-derived oxaloacetate formation and non-oxidative TCA cycle metabolism^49,50^. These findings suggest that cells lacking *Slc25a3* compensate for impaired oxidative metabolism by rerouting carbon through alternative anaplerotic pathways, although this adaptation is insufficient to restore biosynthetic output.

We next considered whether oxidative stress or loss of mitochondrial mass contributes to this phenotype. However, reactive oxygen species levels were modestly reduced in cells lacking *Slc25a3* (Extended Data Fig. 5a,b), and antioxidant treatments failed to restore proliferation (Extended Data Fig. 5c). In addition, markers of mitochondrial content, including citrate synthase and VDAC, were unchanged^51,52^ (Extended Data Fig. 5d). These results indicate that the metabolic defects associated with *Slc25a3* loss do not arise from oxidative stress or reduced mitochondrial abundance.

The accumulation of pyruvate and α-KG suggested that the committed steps of the TCA cycle, catalyzed by the pyruvate dehydrogenase (PDH) and 2-oxoglutarate dehydrogenase (OGDH) complexes, may be impaired. Consistent with this, enzymatic assays revealed reduced PDH and OGDH activity upon loss of *Slc25a3*, both of which were restored by re-expression of WT or mutant SLC25A3 (Fig. 2o,p). These findings identify PDH and OGDH as key targets of mitochondrial Cu availability^53,54^.

Consistent with impaired PDH activity, PDHK1 levels and inhibitory phosphorylation of PDH (Ser293) were increased upon loss of *Slc25a3* (Extended Data Fig. 5e), suggesting a compensatory response to metabolic stress^55,56^. However, this does not account for the reduction in OGDH activity, indicating that mitochondrial Cu loss disrupts TCA cycle function through additional mechanisms. Together, these data demonstrate that loss of mitochondrial Cu impairs oxidative TCA cycle metabolism by disrupting PDH and OGDH activity, leading to metabolic rewiring and reliance on compensatory anaplerotic pathways.

### PC supports survival in low mitochondrial Cu conditions

Given the impairment of PDH- and OGDH-dependent production of TCA cycle metabolites (Fig. 2q,r), we next examined alternative anaplerotic inputs into the TCA cycle. Glutamine-derived carbon entry was assessed using ¹³C-glutamine tracing^57,58^ (Fig. 3a). Loss of *Slc25a3* reduced incorporation of glutamine-derived carbon into α-KG and downstream TCA cycle intermediates (m+5 isotopologues) (Fig. 3b-f), indicating that glutamine anaplerosis is diminished under these conditions.

**Figure 3.**
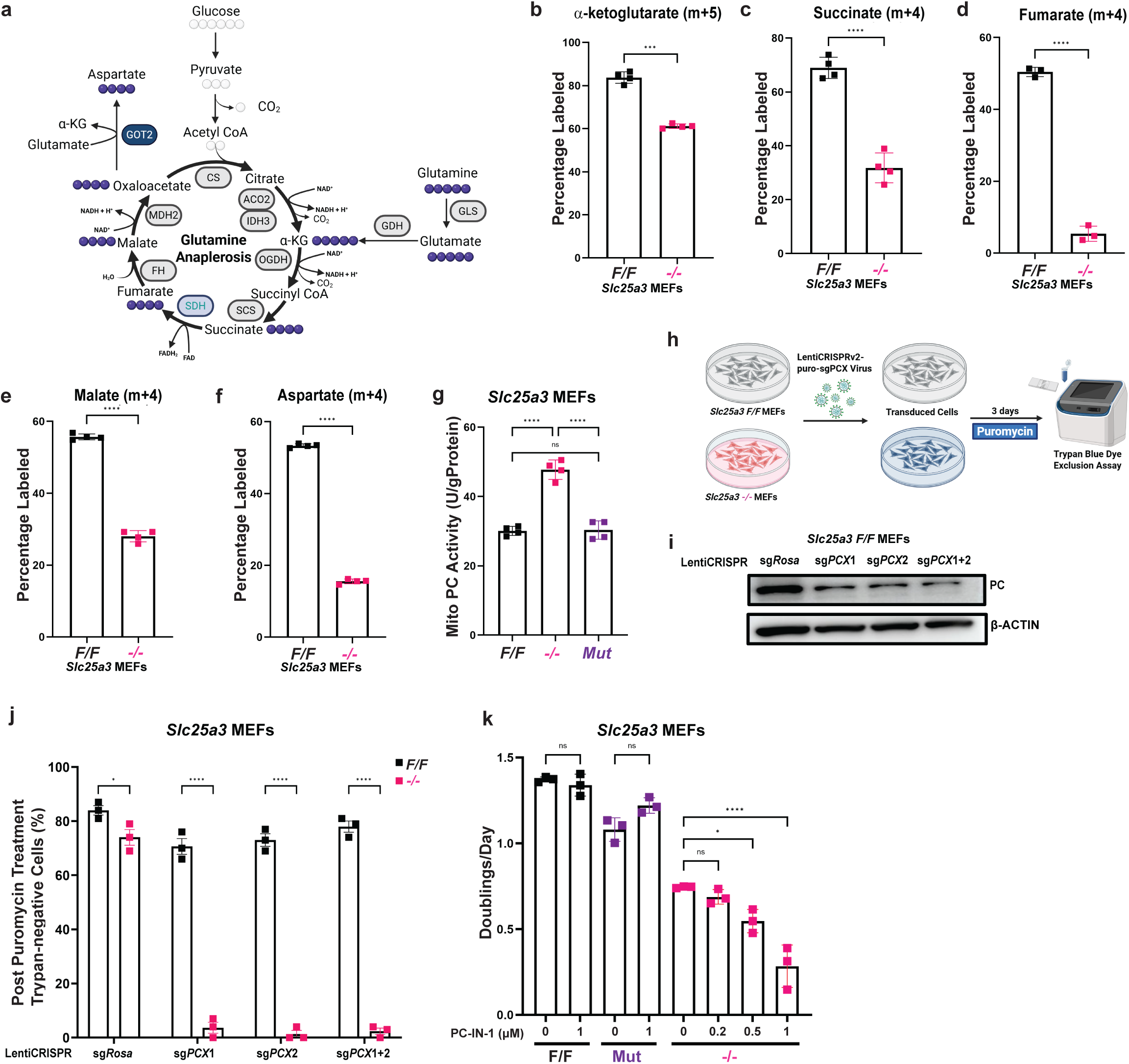
Pyruvate carboxylase is essential for survival in the absence of mitochondrial copper. **a,** Schematic of ^13^C-glutamine tracing pathway from glutamine to glutamine which enters the TCA cycle into α-ketoglutarate through glutamate dehydrogenase (GDH) and cycles in the clockwise direction to produce mostly m+4 states of TCA metabolites into aspartate. **b-f,** ^13^C-glutamine (4mM) stable isotope tracing (6 hour pulse) labeling into **(b)** m+5 α-ketoglutarate, **(c)** m+4 succinate, **(d)** m+4 fumarate, **(e)** m+4 malate, **(f)** m+4 aspartate, in extracts from *Slc25a3 F/F* and *-/-* MEFs, as measured by GC-MS. **g,** Pyruvate Carboxylase activity measured using an activity assay kit in isolated mitochondria from *Slc25a3 F/F*, *-/-* and *Mut* MEFs, normalized to respective protein concentrations. **h,** Schematic workflow representing cells transduced with LentiCRISPRv2 plasmid encoding single guide RNAs (sgRNA) for pyruvate carboxylase (PC), selected with puromycin (1.5µg/mL) for 3 days and subsequently analyzed for cell viability using a trypan blue-dye exclusion assay. **i,** Immunoblot detection of pyruvate carboxylase or β-ACTIN in *Slc25a3 F/F* MEFs transduced with LentiCRISPrv2 plasmid with single guides targeting the Rosa26 locus, or single guides targeting pyruvate carboxylase used separately or in a pool. **j,** Percentage of trypan-negative cells representing cell death post puromycin selection in *Slc25a3 F/F* or *-/-*MEFs, each transduced with either sg*Rosa* virus, or sg*Pcx* (pyruvate carboxylase mouse gene) virus used separately or in a pool. **k,** Doublings per day calculated from cell counts measured using trypan blue dye exclusion in *Slc25a3 F/F* MEFs treated with vehicle or 1µM PC-IN-1, *Slc25a3 Mut* MEFs treated with vehicle or 1µM PC-IN-1, and *Slc25a3 -/-* MEFs treated with vehicle, 0.2µM, 0.5µM or 1µM PC-IN-1. For **g**, **j** and **k**, n=3 biologically independent experiments. For **b-g**, **j** and **k** data is represented as mean ± SEM. ns= not significant, *p≤ 0.05, **p≤0.01, ***p≤ 0.001, ****p≤ 0.0001.

We therefore asked whether pyruvate-derived anaplerosis compensates for impaired oxidative metabolism. Measurement of PC activity in isolated mitochondria revealed increased activity upon loss of *Slc25a3*, which was normalized by re-expression of the mutant transporter (Fig. 3g), indicating that PC is activated in response to mitochondrial Cu depletion^59,60^. To determine whether this increase in PC activity is required for proliferation, we targeted *Pcx* (Fig. 3h,i) and measured cell viability (Fig. 3h). Reduced PC protein levels (Fig. 3i,j) resulted in loss of viability specifically in cells lacking *Slc25a3*, whereas control cells were unaffected (Fig. 3j). Since we were unable to generate *Pcx* knockouts in the *Slc25a3^-/-^* MEFs, we also tested whether PC function is required for proliferation using the small-molecule inhibitor PC-IN-1^61^. PC inhibition had minimal effects on control or mutant-rescued cells but reduced proliferation of cells lacking *Slc25a3* in a dose-dependent manner (Fig. 3k), indicating that these cells depend on PC activity for growth. These findings demonstrate a synthetic dependency on PC upon mitochondrial Cu depletion. Together, these results indicate that loss of mitochondrial Cu impairs oxidative TCA cycle metabolism and creates a reliance on PC-mediated anaplerosis to support cell survival.

### Mitochondrial Cu is required for the stability of lipoylated proteins and the lipoylation machinery

The reliance on PC suggested that the defect in TCA cycle metabolism arises upstream of anaplerotic entry, prompting us to examine the integrity of the PDH and OGDH complexes. Both complexes are multi-enzyme assemblies composed of catalytic E1 subunits that decarboxylate substrates, E2 subunits that carry a covalently attached lipoyl group required for acyl transfer, and a shared E3 subunit (DLD) that regenerates the oxidized cofactor^53,54^. The lipoyl group on the E2 subunit is essential for substrate channeling between catalytic sites and for overall complex activity^62,63^. Immunoblot analysis revealed no change in the catalytic E1 subunits or the shared E3 subunit DLD (Fig. 4a,b). In contrast, the lipoylated E2 subunits DLAT and DLST were nearly undetectable upon loss of *Slc25a3* and restored by re-expression of the mutant transporter (Fig. 4a,b). Rotenone treatment did not recapitulate this effect, indicating that the loss of lipoylated proteins is not a general consequence of ETC inhibition but is specific to mitochondrial Cu depletion (Fig. 4a,b). Other lipoylated proteins, including PDHX1 and DBT, were less affected (Extended Data Fig. 6a), indicating differential sensitivity among lipoylation targets.

**Figure 4.**
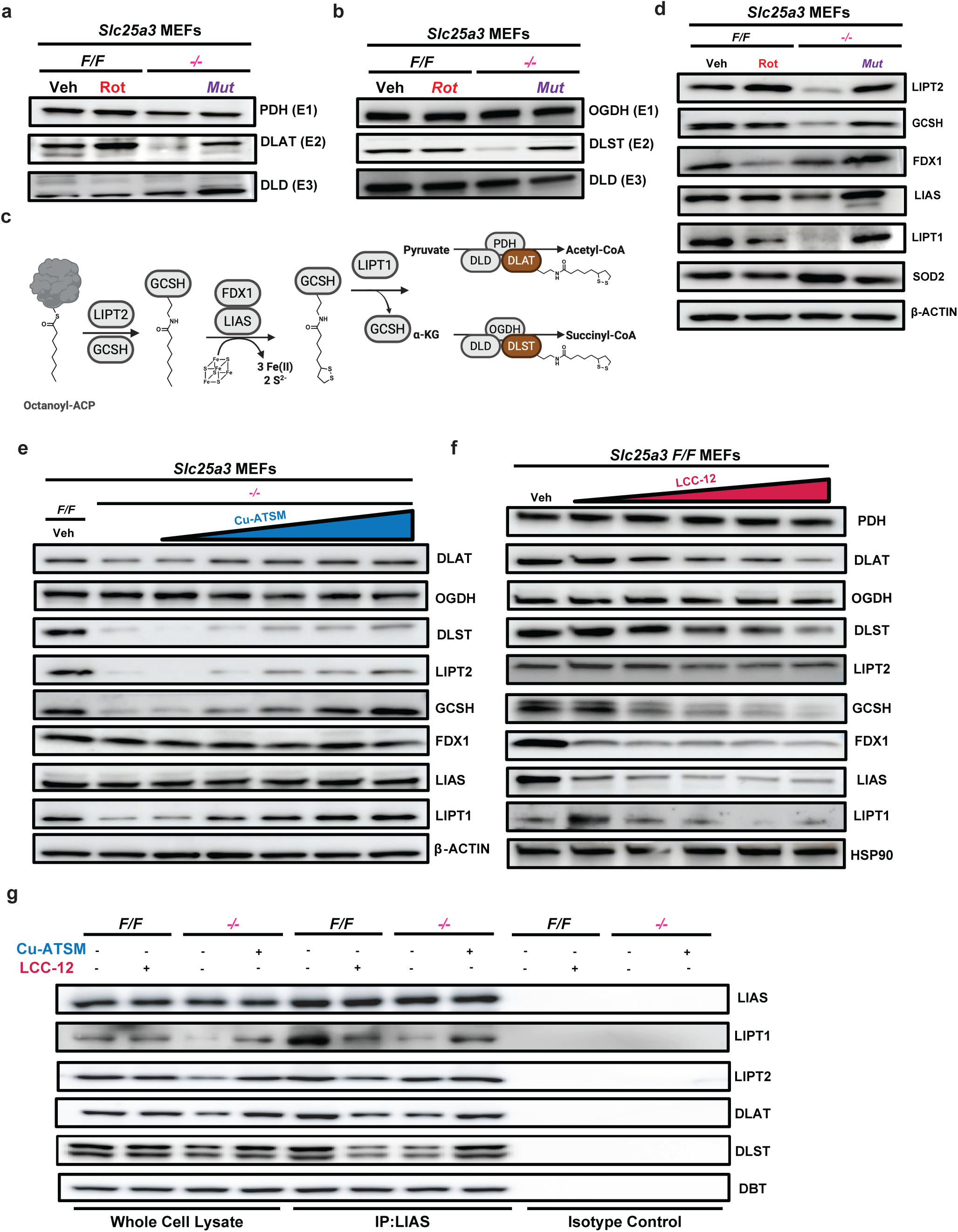
Mitochondrial copper is required for the stability of lipoylated proteins and the lipoylation machinery. **a, b,** Immunoblot detection of **(a)** components of the pyruvate dehydrogenase complex, PDH(E1 subunit), DLAT (E2 subunit), or DLD (E3 subunit), and **(b)** components of the 2-oxoglutarate dehydrogenase complex, OGDH (E1 subunit), DLST (E2 subunit), or DLD (E3 subunit), in *Slc25a3 F/F* MEFs treated with either vehicle or 200nM Rotenone, *Slc25a3 -/-*, *Mut* MEFs. **c,** Schematic representation of the lipoylation pathway where LIPT2 catalyzes the first step of the pathway by transferring an octanoyl moiety to the carrier protein, GCSH; the octanoyl moiety is transformed into a lipoyl group by the coordinated action of FDX1 and LIAS which is then transferred to its final protein partners, DLAT and DLST, by the transferase activity of LIPT1. **d,** Immunoblot detection of LIPT2, GCSH, FDX1, LIAS LIPT1, SOD2 and β-ACTIN levels in *Slc25a3 F/F* MEFs treated with either vehicle or 200nM Rotenone, *Slc25a3 -/-*, *Mut* MEFs. **e, f,** Immunoblot detection of **(e)** DLAT, OGDH, DLST, LIPT2, GCSH, FDX1, LIAS,LIPT1 andβ-actin levels or **(f)** PDH, DLAT, OGDH, DLST, LIPT2, GCSH, FDX1, LIAS, LIPT1 and HSP90 levels in **(e)** SLC253 *F/F* MEFs, treated with either vehicle or increasing concentrations of LCC-12 (10, 20, 30, 40, 50µM) for 24 hours or **(f)** in *Slc25a3 F/F* MEFs treated with vehicle or *Slc25a3* -/- MEFs treated with either vehicle or increasing concentrations of Cu-ATSM(25, 50, 100, 250, 500nM) for 24 hours. **g,** Immunoblot detection of co-immunoprecipitates of LIAS with LIPT1, LIPT2, DLAT, DLST, and DBT from *Slc25a3 F/F* MEFs treated with either vehicle or LCC-12 (30µM) for 24 hours and *Slc25a3 -/-* MEFs treated with either vehicle or Cu-ATSM (250nM) for 24 hours. For **a**, **b**, **d** and **e-f**, n=3 biologically independent experiments.

We next examined components of the lipoylation pathway (Fig. 4c). Loss of *Slc25a3* reduced LIPT2, GCSH and LIPT1, whereas FDX1 and LIAS levels were largely unchanged (Fig. 4d). Expression of the Cu-transport-competent *Slc25a3* mutant restored these protein levels, linking the phenotype directly to mitochondrial Cu availability. Other mitochondrial matrix proteins like SOD2 were not affected signifying that this is specific to certain proteins involved in protein lipoylation. Although transcript levels of these components were also decreased (Extended Data Fig. 6b), the magnitude of this reduction was modest relative to the pronounced loss of the corresponding proteins, indicating that transcriptional changes alone cannot account for the phenotype^64–66^.

To distinguish between transcriptional and post-transcriptional mechanisms, we used the lipophilic compound, Cu-ATSM, to bypass SLC25A3-dependent mitochondrial Cu import^67,68^. In cells with a compromised ETC, Cu-ATSM utilizes the enhanced reductive environment to convert its central Cu(II) ion to its Cu(I) state and releases it into the mitochondrial matrix. Cu-ATSM treatment did not restore mRNA abundance (Extended Data Fig. 6c) but resulted in a dose-dependent recovery of DLAT, DLST, LIPT2, GCSH, and LIPT1 protein levels, whereas FDX1, LIAS, and OGDH remained largely unchanged (Fig. 4e). The rapid restoration of protein levels in the absence of transcriptional recovery supports a post-transcriptional mechanism. Together, these data indicate that mitochondrial Cu is sufficient to stabilize lipoylated proteins and key components of the lipoylation machinery. The discordance between mRNA and protein changes further raises the possibility that the observed reduction in transcript levels reflects a secondary, phosphate-driven phenotype rather than a primary regulatory effect.

To test whether these effects are directly driven by mitochondrial Cu availability, we depleted mitochondrial Cu pharmacologically using LCC-12^69^. This treatment recapitulated the loss of DLAT and DLST, as well as LIPT2, GCSH and LIPT1, in a dose-dependent manner (Fig. 4f). Notably, FDX1 and LIAS were reduced upon Cu chelation. Given that LIAS is an iron-sulfur (Fe-S) cluster enzyme^64,70^, this observation raises the possibility that acute mitochondrial Cu depletion perturbs Fe-S cluster stability or function, although this does not fully account for the selective loss of lipoylated proteins.

Finally, we next examined whether Cu influences the integrity of a previous undescribed lipoylation machinery complex. Co-immunoprecipitation of LIAS revealed association with LIPT1, DLAT, and DLST under control conditions, which was disrupted upon Cu depletion and partially restored by Cu-ATSM treatment of *Slc25a3^-/-^*cells (Fig. 4g). These findings suggest that mitochondrial Cu supports the stability of a multiprotein lipoylation complex, consistent with a post-translational mechanism, and provides a potential explanation for the coordinated destabilization of the pathway.

### Cu directly binds to the lipoyl groups of mitochondrial proteins

Although we established that lipoylated E2 proteins and components of the lipoylation machinery are destabilized in a Cu-dependent manner, these studies did not assess whether the E2 subunits remain lipoylated in cells lacking *Slc25a3*. Immunoblot analysis using a lipoic acid-specific antibody revealed a clear reduction in lipoylated DLAT and DLST in cells deficient in mitochondrial Cu, which was restored upon re-expression of a Cu-transport-competent *Slc25a3* mutant^20,71^ (Fig. 5a). To test this using an orthogonal approach, we first employed a butyraldehyde-based chemical probe (BAP), a thiol-reactive reagent that selectively reacts with reduced lipoyl groups in the presence of TCEP^72–74^ (Fig. 5b). In cell lysates, BAP reactivity was detected by in-gel fluorescence following click chemistry, and was enhanced by TCEP treatment, consistent with increased accessibility of reduced lipoyl groups under these conditions (Extended Data Fig. 7a). BAP labeling was diminished in cells depleted of mitochondrial Cu either pharmacologically (LCC-12) or genetically (loss of *Slc25a3*) which correlates with the absence of the proteins altogether and was partially restored by Cu-ATSM treatment (Extended Data Fig. 7a). Notably, the reduction in BAP signal closely paralleled the loss of DLAT and DLST protein abundance, indicating that mitochondrial Cu primarily affects protein stability rather than directly impairing lipoylation of existing proteins. Consistent with this, enrichment of BAP-reacted proteins followed by immunoblotting showed reduced recovery of DLAT and DLST upon Cu depletion, which was restored by Cu repletion, whereas DBT was largely unaffected (Fig. 5c), indicating differential sensitivity among lipoylated proteins as observed in Extended Data Fig. 6a.

**Figure 5.**
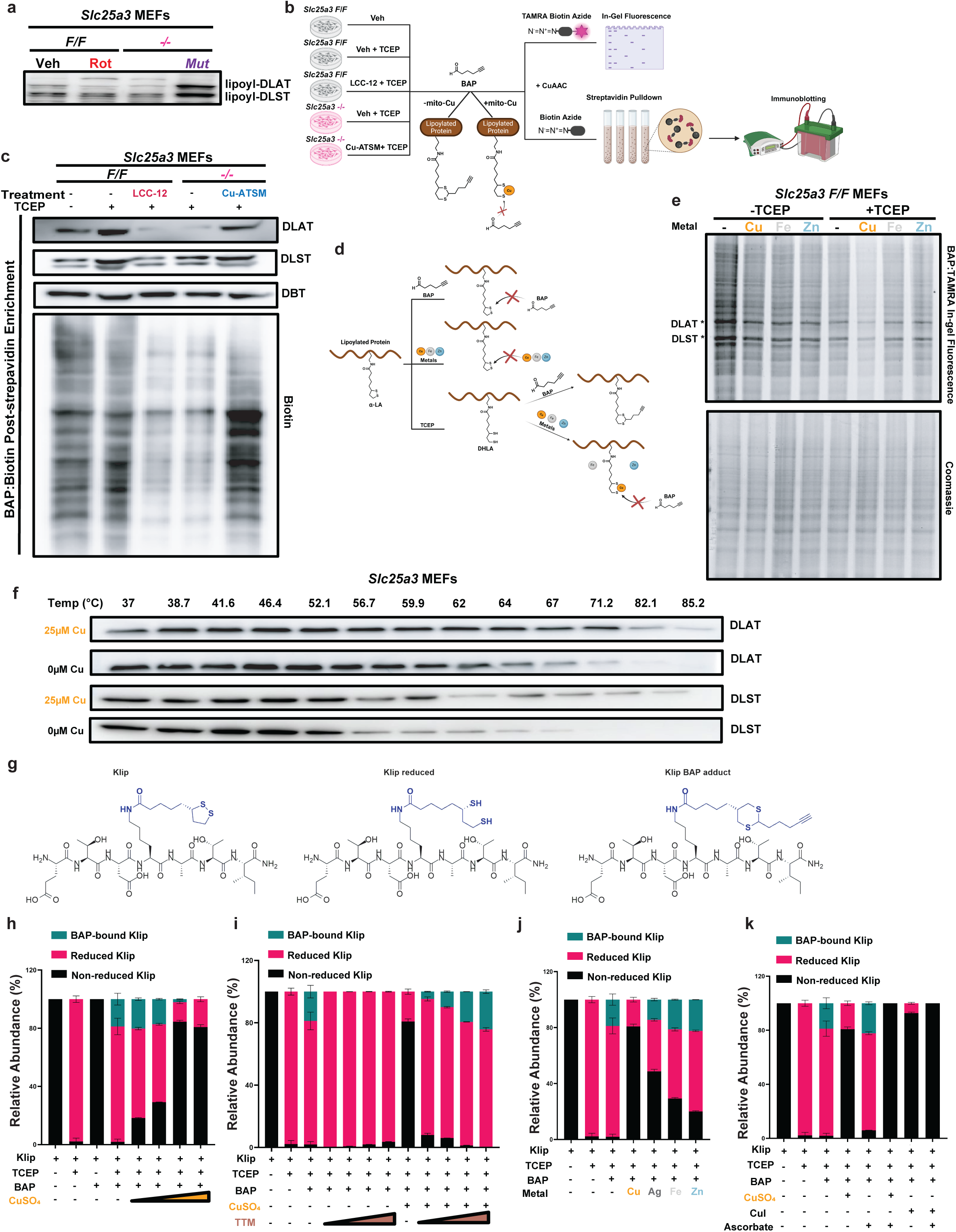
Copper directly binds to the lipoyl tails of mitochondrial proteins. **a,** Immunoblot detection of lipoic acid in *Slc25a3 F/F* MEFs treated with either vehicle or 200nM Rotenone, *Slc25a3 -/-*, *Mut* MEFs. **b,** Schematic representation of BAP binding to lipoic acid tails once reduced with TCEP, forming a cyclic BAP conjugate. Copper competition with BAP binding at the lipoyl tails. BAP treated lysates subjected to click reactions and subsequent streptavidin pulldown for immunoblot detection of target proteins. **c,** Immuno-blotting of post-streptavidin enrichment from BAP labeling of DLAT, DLST, DBT, and Biotin in *Slc25a3 F/F* MEFs treated with vehicle, lysates treated with TCEP (2mM) for 1 hour, or LCC-12 (30µM) for 24 hours and those lysates treated with TCEP (2mM), and *Slc25a3 -/-* MEFs treated with either vehicle or Cu-ATSM (250nM) for 24 hours and those lysates treated with TCEP (2mM) for 1 hour. **d,** Schematic representation of lipoyated proteins with either oxidized lipoyl tails, or the lipoyl tails reduced with TCEP, binding BAP and competing for binding the reduced lipoyl tails with different metals (Cu, Zn, Fe)**. e,** In-gel fluorescence imaging and Coomassie staining of *Slc25a3 F/F* MEF lysates treated with or without TCEP (2mM) for 1 hour, then either with ddH_2_O or CuSO_4_, FeCl_3_, or ZnCl_2_,(25µM) for 1 hour and then BAP (2mM) for 12 hours. **f,** Cellular thermal shift assay investigating changes in stability displayed by DLAT and DLST in the presence or absence of 25µM CuSO_4_, heated for 6 minutes over a temperature range of 37-85.2 □, in *Slc25a3 F/F* lysates. **g,** Schematic representing the synthetic peptide synthesized from the DLAT sequence (246-252, with K249 lipoylated with α-lipoic acid) in its oxidized form (cyclic disulfide form), its reduced form (dihydrolipoic acid form) and its BAP-bound conjugate form. **h-k,** Relative abundance of the following species: DLAT(246-252)K249lip, retention time ∼7.05 min, [M+H]+ 964.44; DLAT(246-252)K249lip reduced, retention time ∼7.14 min, [M+H]+ 966.46 and DLAT(246-252)K249lip BAP conjugate, retention time ∼8.14 min, [M+H]+ 1044.50 detected using MS in reactions containing **(h)** Klip only, Klip reduced with TCEP, Klip and BAP, Klip reduced with TCEP, BAP and increasing concentrations of CuSO_4_ (2.5, 5, 10, 25µM) for 4 hours, **(i)** Klip only, Klip reduced with TCEP, Klip reduced with TCEP, and BAP, Klip reduced with TCEP, BAP and increasing concentrations of TTM (0.25, 1, 2.5, 5µM), Klip reduced with TCEP, BAP and CuSO_4_ (25µM), Klip reduced with TCEP, BAP, CuSO_4_ (25µM)with increasing concentrations of TTM (0.25, 1, 2.5, 5µM), **(j)** Klip only, Klip reduced with TCEP, Klip reduced with TCEP, and BAP with CuSO_4_ (25µM), AgNO_3_ (25µM), FeCl_3_ (25µM) or ZnCl_2_ (25µM) for 4 hours, or **(k)** Klip only, Klip reduced with TCEP, Klip reduced with TCEP and BAP, Klip reduced with TCEP, BAP and either CuSO_4_ (25µM), sodium ascorbate (400µM), both CuSO_4_ and sodium ascorbate, CuI (25µM), or both CuI and sodium ascorbate, for 4 hours. For **a, c, e** and **f** n=3 independent biological experiments. For **h-k** n=3 independent replicates. For **h-k** data represented as mean ± SEM.

To directly test whether Cu interacts with lipoyl groups, we examined whether Cu could compete with BAP reactivity in reduced lysates (Fig. 5d). Addition of Cu, but not other divalent metals, decreased BAP labeling, indicating that Cu binds to the same thiol groups targeted by the probe (Fig. 5e). In agreement, we next assessed whether Cu binding stabilizes lipoylated proteins. Cellular thermal shift assays revealed that Cu supplementation increased the thermal stability of DLAT and DLST (Fig. 5f) consistent with stabilization of their tertiary fold upon Cu binding^75,76^.

To further define this interaction, we used a synthetic lipoylated peptide (Klip) (Fig. 5g). Mass spectrometry analysis showed that reduction of the lipoyl group enabled BAP labeling, whereas addition of Cu reduced BAP reactivity and increased the oxidized species (Fig. 5h), indicating that Cu interferes with probe access to reduced lipoyl thiols. This effect is progressively reversed by increasing concentrations of the Cu chelator TTM, which restores BAP labeling in a dose-dependent manner^77^ (Fig. 5i). These results demonstrate that Cu directly inhibits probe access to lipoyl thiols and that this interaction is reversible upon Cu chelation. Comparison with other transition metals revealed that iron, zinc and silver promoted oxidation but did not block BAP labeling (Fig. 5j), whereas Cu both promoted oxidation and inhibited probe binding (Fig. 5j), consistent with direct Cu coordination to lipoyl thiols or Cu-mediated oxidation of the reduced lipoyl group^78,79^. In the absence of BAP, TCEP treatment shifted Klip predominantly into the reduced form. Addition of CuSO_4_ progressively reversed this effect, increasing detection of non-reduced Klip in a dose-dependent manner despite the presence of TCEP (Extended Data Fig. 7b). This finding suggests that Cu either promotes re-oxidation of the reduced lipoyl group or coordinates the reduced lipoyl thiols in a manner that prevents their detection as free reduced species. Finally, both CuSO_4_ and Cu(I) reduced BAP labeling, and reduction of Cu^2+^ to Cu⁺ enhanced this effect (Fig. 5k), indicating that Cu⁺ is the active species^80^. Together, these findings demonstrate that mitochondrial Cu directly binds to reduced lipoyl groups in mitochondrial proteins, stabilizing lipoylated enzymes and providing a mechanistic basis for impaired PDH and OGDH activity upon Cu depletion.

### Pharmacological restoration of mitochondrial Cu rescues mTORC1 signaling and TCA cycle activity

Having established that Cu directly engages lipoyl groups (Fig. 5e-k), we next tested whether restoration of lipoylated protein stability would rescue downstream metabolic and signaling defects. Cu-ATSM treatment increased phosphorylation of ULK1, p70S6K and 4E-BP1 in a dose-dependent manner (Fig. 6a), indicating recovery of mTORC1 activity. Consistent with restoration of anabolic signaling, Cu-ATSM improved proliferation of cells lacking *Slc25a3* in a dose-dependent manner (Fig. 6b), although growth was not fully restored to control levels.

**Figure 6.**
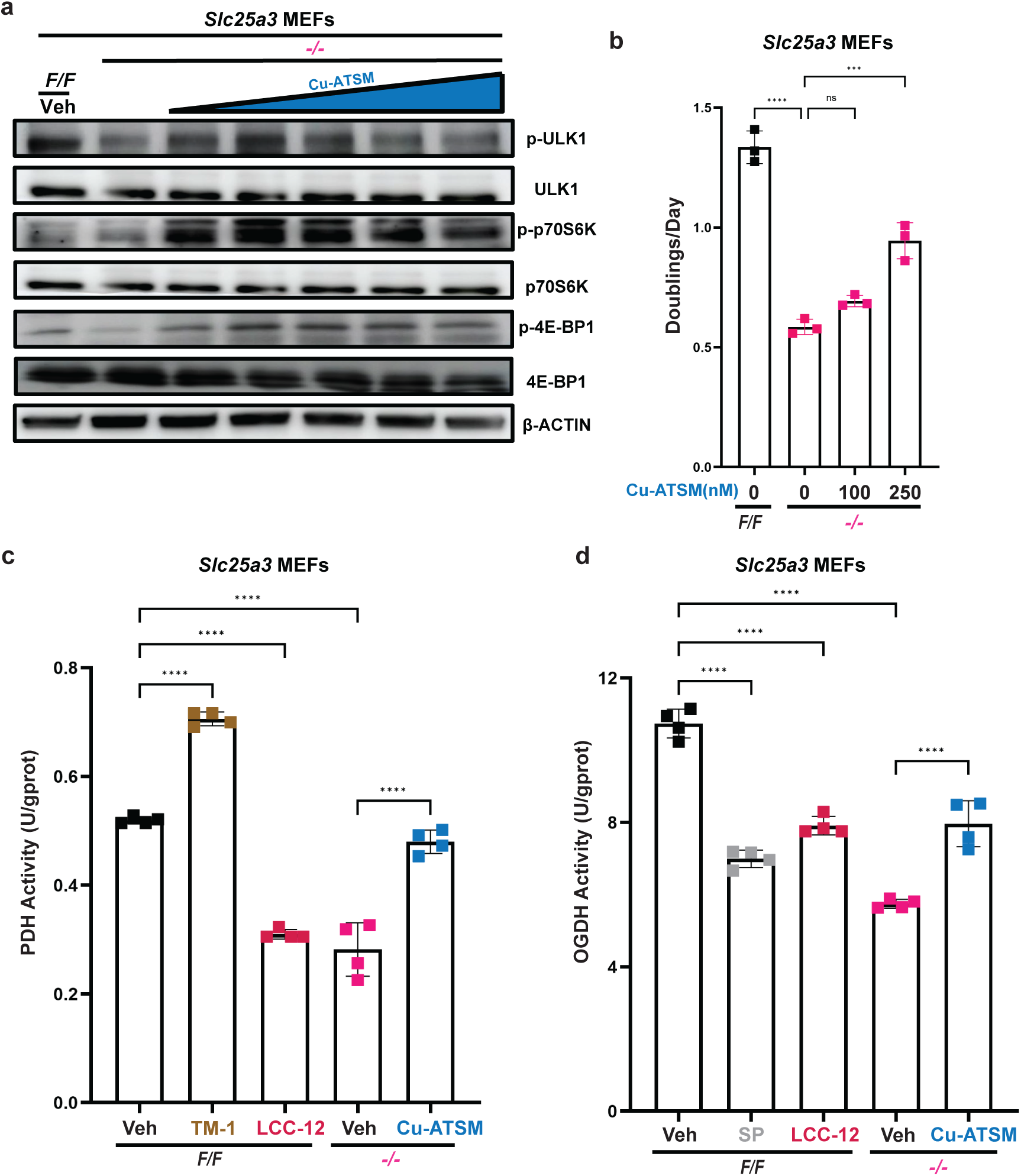
Pharmacological restoration of mitochondrial copper rescues TCA cycle activity and mTORC1 signaling. **a,** Immunoblot detection of p-ULK1, total ULK1, p-p70S6K, total p70S6K, p-4E-BP1, total 4E-BP1 and β-ACTIN in *Slc25a3 F/F* MEFs treated with vehicle or *Slc25a3 -/-* MEFs treated with either vehicle or increasing concentrations of Cu-ATSM(25, 50, 100, 250, 500nM) for 24 hours. **b,** Doublings per day calculated from cell counts measured using trypan blue dye exclusion in *Slc25a3 F/F* MEFs treated with vehicle or *Slc25a3 -/-* MEFs treated with either vehicle or increasing concentrations of Cu-ATSM (100, 250nM) for 72 hours. **c,** Pyruvate Dehydrogenase activity measured using an activity assay kit in isolated mitochondria from *Slc25a3 F/F* MEFs treated with vehicle, TM-1 (10µM) or LCC-12 (30µM) for 24 hours, or *Slc25a3 -/-* MEFs treated with either vehicle or Cu-ATSM (250nM) for 24 hours, normalized to respective protein concentrations. **d,** 2-Oxoglutarate Dehydrogenase activity measured using an activity assay kit in isolated mitochondria from *Slc25a3 F/F* MEFs treated with either vehicle, succinyl phosphonate (100µM) or LCC-12 (30µM), or *Slc25a3 -/-* MEFs treated with either vehicle or Cu-ATSM (250nM) for 24 hours, normalized to respective protein concentrations. For **a-d** n=3 biologically independent experiments. For **b-d** data is represented as mean ± SEM. ns= not significant, ***p≤ 0.001, ****p≤ 0.0001.

We next tested whether this rescue extends to enzymatic activity within the TCA cycle. PDH and OGDH activities were reduced upon loss of *Slc25a3* and following mitochondrial Cu chelation with LCC-12 (Fig. 6c,d). Cu-ATSM treatment restored both PDH and OGDH activity, indicating recovery of oxidative TCA cycle function. Succinyl phosphonate, the competitive OGDH inhibitor, and TM-1, the PDHK1 inhibitor, which serves to relieve the inhibition on the PDH complex, worked as expected. Together, these results demonstrate that mitochondrial Cu is sufficient to restore lipoylated proteins, PDH and OGDH activity and mTORC1-dependent growth signaling, linking Cu-dependent stabilization of lipoyl groups to metabolic and proliferative control. These findings are consistent with a model in which loss of mitochondrial Cu destabilizes lipoylated proteins, leading to their degradation and consequent impairment of TCA cycle function.

## Discussion

This study identifies a central role for mitochondrial Cu in maintaining the stability and function of lipoylated TCA cycle enzymes. Loss of the mitochondrial Cu transporter *Slc25a3* results in the selective depletion of lipoylated proteins, including the E2 subunits of the PDH and OGDH complexes (DLAT and DLST), as well as key components of the lipoylation machinery. This effect is not recapitulated by acute inhibition of the ETC, indicating that it is not a general consequence of respiratory dysfunction but instead reflects a specific requirement for mitochondrial Cu. The resulting metabolic phenotype, characterized by impaired oxidative TCA cycle metabolism, aspartate depletion, suppression of mTORC1 signaling, and dependence on PC positions Cu as a critical regulator of mitochondrial metabolism.

Cu has long been understood as a catalytic cofactor required for Complex IV or COX activity^1,2^. However, we observe that loss of mitochondrial Cu leads to reduced abundance of subunits across multiple ETC complexes, suggesting that the consequences of Cu depletion extend beyond Complex IV. This phenotype differs from classical models of systemic Cu deficiency, where defects are largely restricted to Complex IV^81,82^. While the precise mechanism underlying this broader ETC destabilization remains unclear, our data indicate that it is unlikely to be driven by generalized oxidative stress or loss of mitochondrial mass. Instead, these findings suggest that mitochondrial Cu may influence the stability or assembly of multiple protein complexes, either directly or through its effects on TCA cycle function.

Importantly, our data demonstrate that restoring ETC activity or redox balance is insufficient to rescue the proliferative defect caused by *Slc25a3* loss. This indicates that the primary metabolic lesion lies upstream of the ETC, at the level of TCA cycle enzyme activity. Specifically, loss of PDH and OGDH activity prevents the generation of key metabolic intermediates required for biosynthesis, explaining why restoring electron flow alone does not normalize growth^15,16,39^. These findings establish that Cu-dependent stabilization of lipoylated proteins is a key determinant of metabolic capacity.

The PDH and OGDH complexes are tightly regulated enzymes that control entry into, and progression through, the TCA cycle^53,54^. While PDH activity is classically regulated by phosphorylation and OGDH by allosteric inputs, our data reveal an additional mode of regulation in which Cu availability governs the stability of the lipoylated E2 subunits required for complex function. In the absence of Cu, DLAT and DLST are selectively lost, leading to impaired enzyme activity and collapse of oxidative metabolism. This identifies Cu as a structural regulator of these complexes, extending its role beyond that of a catalytic cofactor.

A major consequence of this defect is depletion of aspartate, a key metabolite linking mitochondrial function to nucleotide biosynthesis and mTORC1 signaling^15,17^. Loss of aspartate limits anabolic growth and activates the integrated stress response, contributing to the observed proliferative defect^18,83^. In parallel, reduced aspartate levels may relieve inhibition of PC, promoting compensatory anaplerotic metabolism^59,84^. While this adaptation partially restores TCA cycle intermediates, it is insufficient to fully support biosynthetic demands, resulting in a continued dependence on PC activity for survival.

The selective loss of lipoylated proteins and lipoylation machinery components raises the question of how these proteins are destabilized. One possibility is that the absence of Cu renders lipoylated proteins susceptible to post-translational modifications that target them for degradation. Previous studies have shown that components of the lipoylation pathway can be regulated by the ubiquitin-proteasome system, and that thiol modifications such as S-glutathionylation can promote protein turnover^85–87^. While the specific mechanisms responsible for DLAT and DLST degradation remain to be defined, our data are consistent with a model in which Cu stabilizes these proteins and protects them from degradation.

Our biochemical and chemoproteomic analyses provide direct evidence that Cu binds to lipoyl groups. We propose that Cu(I) interacts with the reduced thiol groups of the lipoyl moiety, stabilizing its conformation and preventing aberrant oxidation or modification^78,80^. In the absence of Cu, these thiols may become exposed to oxidative or covalent modifications that promote protein destabilization. This model provides a mechanistic explanation for the coordinated loss of lipoylated proteins and associated enzymes, which likely exist within a multiprotein complex^65,66^.

These findings also place our work in conceptual contrast with the recently described process of cuproptosis. In that setting, Cu overload promotes aggregation of lipoylated proteins and induces cell death^20^. Here, we show that Cu depletion leads to loss of the same proteins. Together, these observations suggest that lipoylated proteins act as sensitive reporters of Cu availability, with distinct cellular outcomes depending on Cu levels. Rather than a linear relationship, these data support a biphasic model in which both Cu deficiency and excess disrupt mitochondrial function through effects on lipoylated proteins^6,88^.

Finally, our results have potential therapeutic implications. We show that pharmacological delivery of mitochondrial Cu using Cu-ATSM restores lipoylated protein abundance, PDH and OGDH activity and cell proliferation, suggesting that this approach may be beneficial in diseases associated with mitochondrial Cu deficiency^67,89^. Conversely, mitochondrial Cu depletion using LCC-12 phenocopies the metabolic defects observed upon *Slc25a3* loss, raising the possibility that targeting mitochondrial Cu could be exploited in cancers dependent on oxidative metabolism^69,90^. The observed dependency on PC further suggests potential combination strategies^61,91^.

In summary, this study establishes a model in which mitochondrial Cu regulates central carbon metabolism by stabilizing lipoylated TCA cycle enzymes. Loss of Cu disrupts this system, leading to metabolic collapse and impaired cell growth. These findings expand the role of Cu in biology from a catalytic cofactor to a regulator of protein stability and metabolic function.

Several questions remain. The mechanisms governing degradation of lipoylated proteins in the absence of Cu, the identity of potential quality control pathways involved, and the extent to which Cu regulates additional mitochondrial protein complexes are areas for future investigation. Addressing these questions will further define how Cu homeostasis integrates with mitochondrial function and cellular metabolism^7,8,92^.

## Methods

### Cell Lines

*Slc25a3 Flox/Flox (F/F)* and *-/-* MEFs were a gift from Scot Leary. *Slc25a3 F/F* and *-/-* immortalized MEFs were stably infected with retroviruses derived from pBABE or pWZL (see the ‘Plasmids’ section) or lentiviruses derived from pLentiCRISPRV2 (see the ‘Plasmids’ section) using established protocols.

### Plasmids

pWZLblasti-SLC25A3-WT was created by cloning the human SLC25A3 cDNA sequence with an N terminus FLAG tag into the pWZLblasti-CTR1^WT^ (Addgene plasmid 53157) replacing the CTR1^WT^ sequence. pWZLblasti-SLC25A3-Mut was created by site-directed mutagenesis of the pWZLblasti-SLC25A3-SLC25A3-WT plasmid to change Leucine175 to Alanine. pLentiCRISPRv2-puro (Addgene plasmid 52961) was used to create pLentiCRISPRv2-puro-Rosa26-sgRNA (mouse Rosa26 target sequence 5′-CCCGATCCCCTACCTAGCCG), and the pLentiCRISPRv2-puro-PCX-sgRNA (mouse Pcx target sequences 5’-GACACCGGCCGCATTGAGGT, 5’-GCACGCACGAAACACTCGGA). pCDH-puro-AOX-FLAG was created by cloning the AOX-FLAG sequence from pCW57.1-AOX-FLAG (Addgene plasmid 177984) into the pCDH-puro-cMyc (Addgene plasmid 46970) plasmid using primers to include the FLAG tag and replacing the cMyc sequence. pCDH-puro-NDI1 was created by cloning the NDI1 sequence from pLV-EF1a-NDI1-mRFP1 (gift from Navdeep Chandel) into the pCDH-puro-cMyc plasmid by replacing the cMyc sequence. pCDH-puro-LBNOX and pCDH-puro-mitoLBNOX was created by cloning either the Lbnox or mitoLbNox sequence from pUC57-LbNox (Addgene plasmid 75285) or pUC57-mitoLbNox (Addgene plasmid 74448) into the pCDH-puro-cMyc plasmid using primers to include the FLAG tag and replacing the cMyc sequence.

### Mitochondrial Isolation

Cells from a confluent 15-cm dish was harvested and wash with 1x PBS and re-suspended in RSB Hypo Buffer (10 mM NaCl, 1.5mM MgCl_2_, 10mM Tris-HCl at pH7.5). The cells were allowed to swell on ice for 15 minutes. Swollen cells were transferred to a 5mL Dounce homogenizer and broken open with 20-25 strokes. A 2.5x MS Homogenization buffer was added to the cell suspension, containing intact mitochondria, to make a 1x MS Homogenization buffer solution (210 mM mannitol, 70 mM sucrose, 5 mM Tris-HCl at pH 7.5, 1 mM EDTA at pH 7.5). The cell solution was centrifuged at 1,300g for 5 minutes at 4□. The supernatant was transferred to a new tube and centrifuged twice more in sequence to remove the nuclear fraction and other cellular debris. The supernatant was then transferred to a new tube and centrifuged at 15,000g for 15 minutes at 4□. The mitochondrial pellet was washed once with 1x MS Homogenization buffer. The mitochondrial pellet was re-suspended in Extraction Buffer (150 mM sodium acetate, 30 mM HEPES, 1mM EDTA, 12% glycerol and pH 7.5) and stored at -80□ till required.

### Cytochrome Oxidase Activity

Mitochondrial pellets from the appropriate cells with the indicated treatments were processed for their cytochrome oxidase activity using the Abcam Cytochome C Oxidase Activity Assay Kit (Abcam ab239711) according to manufacturer’s instructions. The activities were normalized to mitochondrial pellet protein concentrations calculated by a BCA assay.

### Oxygen Consumption Rate (OCR)

40,000 cells of each cell line were plated in Seahorse XF Pro 96-well plates (Agilent 103734-100) and OCR was measured using the Seahorse XF Pro Analyzer using the Seahorse XF Cell Mito Stress Test Kit (Agilent 103015-100) with sequential addition of 1.5µM oligomycin, 1µM FCCP and 500nM Rotenone. Data was analyzed using Seahorse Analytics software.

### NAD/NADH Ratio Measurement

Mitochondria from SLC25A3 F/F, -/- and Mut MEFs were isolated accordingly to the protocol above. Mitochondria stored in the Extraction Buffer was centrifuged at 15,000g for 15 minutes and the pellets were processed for their NAD^+^/NADH ratio using the Promega NAD/NADH-Glo Assay (Promega G9071) according to manufacturer’s instructions.

### ATP Measurement

Whole cell pellets or mitochondrial pellets from SLC25A3 F/F, -/-and Mut MEFs were processed for their ATP levels using the ATPlite Luminescence Assay System (Revvity Health Sciences 6016943) according to manufacturer’s instructions.

### Trypan Blue Dye Exclusion Assay

Cell viability after treatment with indicated treatments was measured using a Countess III cell counter (ThermoFisher). Briefly, the appropriate cell lines were plated in 6-well plates. The next day, vehicle or treatment was added for the indicated times. Cells were then harvested and concentrated to 2–5 million cells/ml. Trypan Blue was added immediately before counting for cell viability. Doubling Time was calculated using the formula: Time x ln(2)/ln(final cell number/initial cell number). Doublings per day was calculated using the formula: 24/Doubling Time.

### Quantitative PCR

RNA was purified from MEFs using TRIzol Reagent (Thermo Fisher Scientific 15596026) following standard protocols and reverse transcribed to cDNA using using oligo dT primers (Promega C1101). The cDNA was then quantified using the Taqman probe Mm00433246_m1 to detect mouse Fdx1, Mm00513470_m1 to detect mouse Dlst, Mm03015639_m1 to detect mouse Lipt1, Mm00840330_g1 to detect mouse Lipt2, Mm00522477_m1 to detect mouse Lias, Mm00783118_s1 to detect mouse Gcsh, Mm00455160_m1 to detect mouse Dlat, Mm02619580_g1 to detect mouse Actb, Mm00441480_m1 to detect mouse Glut1, Mm01612132_g1 to detect mouse mouse Ldha using the Applied Biosystems QuantStudio6 Pro Real-Time PCR System. Relative mRNA expression levels were normalized to Actb and analyzed using comparative ΔΔ*C*t method.

### Measurement of Glucose Uptake and Lactate Production

Cells were plated at a density of 25,000 cells/well in 6-well plates in a total of 2 mL of culture media. Wells containing media but no cells were kept as controls for normalization of metabolite concentrations in media. After 24 hours post-seeding, media was removed, wells were washed with 1X dPBS, and fresh media was added to each well. Samples of culture media were collected 48 hours post-media change and stored at −80°C until analysis. For normalization purposes, number of cells per well was determined manually using an automated cell counter 24 hours after seeding and at endpoint. Quantification of glucose and lactate was determined enzymatically with a bioanalyzer (YSI2950, YSI Incorporated, Yellow Springs, OH, USA). Rate of metabolite uptake/production was calculated by subtracting the concentration of glucose/lactate in control media and dividing by the cell number area under the curve.

### Immunoblot analysis

The indicated cell lines with appropriate conditions were washed with cold phosphate-buffered saline (PBS) and lysed with ice-cold RIPA buffer containing 1× EDTA-free Halt protease and phosphatase inhibitor cocktail halt protease and phosphatase inhibitors (Thermo Fisher Scientific). The protein concentration was determined by BCA protein assay (Pierce) using BSA as a standard. Equal amounts of lysate were resolved by SDS–PAGE using standard techniques, and transferred to a nitrocellulose membrane using a Bio-rad Trans-blot Turbo Transfer System. Protein was detected using the appropriate antibodies in Table 1.

### Surface Sensing of Translation (SUnSET)

The indicated cell lines were plated in 10-cm dishes and subsequently treated with either vehicle or 10 mg/mL puromycin for 15 minutes. The cells were then processed for immunoblot analysis as described above and blotted for puromycin.

### LC-HRMS analysis

Cells were washed with cold 1x PBS and extracted using 80% MeOH (mass-spec grade diluted in water) on ice. Cells were scraped on ice with a cell scraper and centrifuged at 15,000g at 4°C. Metabolite extracts were subjected to LC-HRMS analysis on a Vanquish UHPLC system coupled to a Q Exactive HF Orbitrap mass spectrometer outfitted with a heated electrospray ionization source (Thermo Fisher Scientific). Separations were carried out on an Atlantis Premier BEH Z HILIC VanGuard FIT column (2.1 × 150 mm, 2.5 µm; Waters) at 30°C. Mobile phase A contained 10 mM ammonium carbonate and 0.05% ammonium hydroxide in water, while mobile phase B was acetonitrile. Gradient elution proceeded as follows: 80% B at initial conditions, ramping down to 20% B by 13 minutes, isocratic hold at 20% B to 15 minutes, followed by a return to initial conditions for re-equilibration. The chromatographic flow rate was 150 µL/min with a 5 µL injection volume. Data were acquired in both positive and negative electrospray ionization modes. The MS1 scan range was set to m/z 65–950 for positive mode and m/z 65–975 for negative mode, with a resolving power of 120,000 (at m/z 200) and an AGC target of 3 × 10. Raw files were converted to .cdf format using Xcalibur (v4.0) before import into El-Maven (v0.12.0) for feature extraction and processing. Peaks were extracted at an EIC mass tolerance of ±10 ppm, with metabolite identifications made by comparison of accurate mass and retention time to an in-house authenticated standard library.

### 13C Isotope Tracing

SLC25A3 F/F or -/- MEFs were plated in 6-well plates in triplicate at 100k cells/well. Cells were left to adhere overnight and the next day, media was aspirated off and cells were washed with 1x PBS before basal tracing media with supplements was provided. Tracing medium is a DMEM-based medium without glucose, glutamine or pyruvate. ^12^C glucose (10 mM) and ^12^C glutamine (4 mM) was added to base media along with 10% dialyzed fetal bovine serum (Gemini Bio 100-108) for 24 hours to allow for cells to acclimate to the new media. The next morning, the basal tracing medium was refreshed and 2 hours later, cells were washed twice with dPBS before tracing media with ^13^C-metabolites was added. Unlabeled controls were provided with ^12^C tracing media, while ^13^C metabolite wells were provided with ^12^C_5_ glutamine with ^13^C_6_ Glucose or ^12^C_6_ glucose with ^13^C_5_ glutamine. Concentrations of ^13^C metabolites were equal to ^12^C so that cells do not experience any change in metabolite abundance during the tracing period. Cells were incubated with labeled substrates for 6 hours before they were harvested for extraction to be run on GC-MS/LC-MS. Unlabeled control wells were used to correct labeling for natural abundance of isotopologues.

### Metabolite Extraction for GC-MS

Cells were washed with cold 1x PBS at the experimental end point and extracted using 80% MeOH (mass-spec grade diluted in water) on ice. Cells were scraped on ice using a cell scraper and centrifuged at 15,000g at 4°C. The pellet was saved at -80°C. For extraction normalization, an internal standard (norvaline) was added to each sample supernatant to account for sample loss during prep. Samples were then vortexed for 60 seconds prior to centrifugation at 4°C at 21,000g for 15 minutes. The supernatant was saved and was then spun down overnight to dry using a Speedvac (Thermo Scientific SPD130DLX).

Once dried, metabolite pellets were resuspended in 30 uL pyridine containing 10 ug/uL methoxyamine HCl (Sigma 226904) to form methoxime-TBDMS adducts. The samples were then vortexed for 10 seconds each, twice. They were then heated at 70°C for 15 minutes on a heating block. 70 uL MTBSTFA (Sigma 394882) was added to these samples for derivatization, vortexed and heated at 70 °C for a further 60 minutes. Samples were then spun down at 21,000g for 20 minutes at 4°C, and 40uL of the supernatant was transferred to glass vials with glass inserts and stored at 4°C until they were run and analyzed using a gas chromatographer (Agilent 7890B) with a HP-5MS 5% phenyl methyl Silox column (Agilent) coupled to a mass spectrometer (Agilent 5977A). For stable isotope tracing experiments, an unlabeled control was included for each cell line. Natural isotope abundance was performed using FluxFix^93^.

### Metabolite Extraction for Acyl-CoAs

Acyl-CoAs were analyzed by liquid chromatography-high-resolution mass spectrometry (LC-HRMS) with isotope dilution with short-chain acyl-CoA internal standards (ISTD) generated in yeast as described^94,95^. Briefly, 50 µL of short-chain acyl-CoA internal standard was added to cell suspensions in 10% (w/v) trichloroacetic acid and were sonicated with 5 x 0.5-second pulses at 50% intensity (Fisherbrand™ Sonic Dismembrator Model 120 with Qsonica CL-18 sonicator probe). Lysates were centrifuged at 17,000 g for 10 minutes at 4°C and clarified lysates were transferred to a deep-well 96-well plate for loading in a Tomtec Quadra4 liquid handling workstation. On the liquid handling workstation, lysates were applied to an Oasis HLB 96-well elution plate (30 mg of sorbent per well) pre-conditioned and equilibrated with 1 mL of methanol and 1 mL of water, respectively. After de-salting with 1 mL of water, acyl-CoAs were eluted into a deep-well 96-well plate using 1 mL of 25 mM ammonium acetate in methanol. Eluent was evaporated to dryness under nitrogen gas. The dried samples were resuspended in 50 µL of 5% (w/v) sulfosalicylic acid in water. 5 µL injections of each sample were analyzed via LC-HRMS. Acetyl-CoA, propionyl-CoA, and succinyl-CoA lithium salts as well as 5-sulfosalicylic acid were from Sigma-Aldrich. Optima® LC/MS grade acetonitrile (ACN), formic acid, methanol, and water were purchased from Fisher Scientific. Oasis® HLB 96-well elution plates (30 mg of sorbent) were purchased from Waters (P/N: WAT058951). Acyl-CoAs were measured by their predominant M+H ion, and M-507 fragmentation using Tracefinder 5.1. Correction for natural isotopic abundance was performed using Fluxfix^93^.

### Pyruvate Dehydrogenase Activity

Mitochondrial pellets from the appropriate cells with the indicated treatments were processed for their pyruvate dehydrogenase activity using the Abcam Pyruvate Dehydrogenase Activity Assay Kit (Abcam ab287837) according to manufacturer’s instructions. The activities were normalized to mitochondrial pellet protein concentrations calculated by a BCA assay.

### 2-Oxoglutarate Dehydrogenase Activity

Mitochondrial pellets from the appropriate cells with the indicated treatments were processed for their 2-oxoglutarate dehydrogenase activity using the Elabscience α-Ketoglutarate Dehydrogenase Activity Assay Kit (Elabsceince E-BC-K083-M) according to manufacturer’s instructions. The activities were normalized to mitochondrial pellet protein concentrations calculated by a BCA assay.

### Co-Immunoprecipitation

LIAS Co-IP was performed with 1 mg of *Slc25a3 F/F* lysate using Protein G Dynabeads (Thermo, cat#:10004D) according to the manufacturers protocol, with the antibodies shown in Table 1.

### Hex-5-ynal (BAP)

Synthesis of BAP was carried out according to previously published procedures^74^. Briefly, Dess-Martin Periodinane (3.03g, 6.87 mmol, 1 eq.) was stirred in 20mL dichloromethane under a nitrogen atmosphere at 0 °C. Then hex-5-yn-1-ol (0.6 mL, 674 mg, 6.87 mmol) was added drop-wise and the solution was stirred overnight at room temperature. The solution was filtered over a sand core funnel covered with silica gel and the organic layer was evaporated at 4□. After purification by silica gel chromatography, BAP was obtained as thick white oil (650 mg, 6.77 mmol, 98% yield). Analytical data was in correspondence with previously published results, MS calculated for C_6_H_8_O [M + H]^+^, 97.1; observed 97.4.

### BAP labeling experiments

All reactions were carried out in 50 mM citrate buffer pH 4 at 37 °C for 4 hours in 20 μL reaction volumes. Generally, 10x stocks of all reagents were prepared in 50 mM citrate buffer pH 4, with the exception of BAP which was prepared in DMSO. Reactions were carried out with varying compositions at final concentrations of DLAT(246-252)K249lip (10 μM), TCEP (50 μM), BAP (50 μM), CuSO_4_ (2.5-25 μM), CuI/AgNO_3_/FeCl_3_/ZnCl_2_ (25 μM), TTM (2.5-50 μM), sodium ascorbate (400 μM) and BCS (2.5-400 μM). After completion of the reaction, the reactions were quenched by sequential additions of 2 μL 10 mM EDTA and 2 μL of 1% formic acid in MQ-H_2_O (v/v). LC/MS analyses were conducted on an Agilent 1290 Infinity II UHPLC system coupled with an Agilent 6545XT AdvanceBio LC/Q-TOF system equipped with an Agilent Dual Jet Stream ESI source. The following species were quantified in the analyses: DLAT(246-252)K249lip, retention time ∼7.05 min, [M+H]^+^ 964.44; DLAT(246-252)K249lip reduced, retention time ∼7.14 min, [M+H]^+^ 966.46 and DLAT(246-252)K249lip BAP conjugate, retention time ∼8.14 min, [M+H]^+^ 1044.50.

### Butyraldehyde Probe (BAP) Labeled Sample Preparation

SLC25A3 F/F or -/- MEFs with the indicated treatments were scrapped down in 1x PBS on ice and centrifuged for 5 minutes at 300g to pellet the cells. The pellet was washed twice with ice-cold 1x PBS and then re-suspended in Acid Lysis Buffer (41.1 mM Na_2_HPO_4_, 79.45mM Citric acid, pH 3) at room temperature. The samples were centrifuged at 20,000g for 1 hour at room temperature. Protein concentrations were determined using a Bradford assay and samples concentrations were adjusted to 2 mg/mL. TCEP and BAP probe were added to the samples to a final concentration of 2 mM. The samples were shaken for 12 hours at 37°C. To precipitate the proteins, acetone was added to the samples and incubated at -80°C for 1 hour. The samples were subsequently centrifuged at 10,000g for 10 minutes. The pellets were washed with -80°C methanol twice and then re-suspended in 0.4% SDS/PBS solution before subsequent click chemistry steps.

### Click Chemistry Reaction

The general click reaction mix per 25 µL reaction volume was: 1.5 µL of 1.7 mM Tris-benzyltriazolylmethyl-amine (TBTA, Cayman Chemical Company, catalog #18816, in DMSO), 0.5 µL of 50mM CuSO_4_ (Sigma-Aldrich C8027-500G, in Milli-Q H_2_O), 0.5 µL of 13 mg/mL Tris(2-carboxyethyl)phosphine hydrochloride (TCEP, Sigma-Aldrich C4706-2G, in H_2_O), and 0.5 uL of 100 mM Biotin Azide (Specifications below for specific applications). The master mix reagents are added in the order listed above to each respective sample and incubated at room temperature for one hour. After normalizing protein concentration, the sample was split into 25 µg, 250 µg and 1mg fractions. The 25 µg fraction was click labeled with a TAMRA-Biotin Azide (Vector Laboratories CCT-1048, in DMSO) for in-gel fluorescence. The remaining fractions were click labeled with Azide-PEG3-Biotin conjugate (Sigma-Aldrich 762024, in DMSO) for immunoblotting post-streptavidin pulldown.

### In-Gel Fluorescence

After one hour incubation with click-reaction mix, 5x Sample Buffer (0.25 M Tris pH 6.8 Tris, 10% SDS, 50% Glycerol, 25% β-Mercaptoethanol, 0.5% Bromophenol Blue) was diluted to 1X in the samples. The samples were then heated to 95°C for 5 minutes and then loaded onto a Bio-Rad 4-20% Gradient Gel (TGX 10-Well 4561094, TGX 15-Well 4561096). The gels were run until the dye front ran off. The gels were then imaged for fluorescence at a wavelength of 535 nm on the Cytiva ImageQuant800 (Cytiva 29399484).

### Streptavidin Pulldown

The Streptavidin Magnetic Beads (New England Biolabs S1420S) were prepared by washing three times with a conjugation buffer (PBS + 500 mM NaCl), maintaining the beads in a 1:1 slurry. Per each reaction, the biotinylated sample was then mixed with 1 mL of conjugation buffer and 100 µL of the washed 1:1 Streptavidin bead slurry. The samples were then rotated for two hours at room temperature. After rotating, the samples were washed nine times in total: three times with a wash buffer (PBS + 500 mM NaCl + 0.1% Tween 20), three times with PBS, and three times with MilliQ water.

### Cellular Thermal Shift Assay

Whole cell lysate from SLC25A3 F/F MEFs was obtained using RIPA buffer and centrifuged at 12,000g for 5 minutes to remove cellular debris. The lysate was normalized to 2 mg/ML and treated with either water or 25 µM CuSO_4_ and incubated for 1 hour. The lysate from both treatment groups was then heated along a temperature gradient of 37-87□ for 6 minutes. Heat-treated lysates were centrifuged at 21,000g for 30 minutes at 4 ℃. Equal amounts of the supernatant were loaded on to a 10% SDS-PAGE gel and run in 1x Tris-Glycine-SDS buffer. The gel was transferred and immunoblotted according to the steps described above.

### DLAT(246-252)K249lip

The peptide was synthesized by manual Fmoc solid phase peptide chemistry on Rink-Amide MBHA resin. The resin underwent iterative cycles of Fmoc deprotection (20% piperidine in DMF, 30 minutes), amino acid coupling (4 eq. Fmoc AA, 4 eq. HATU, 8eq. DIPEA in DMF, 1 hour) and washing (3 × DMF) until completion of the sequence. At the K249 position, ivDde-protected lysine was introduced. Before final Fmoc-deprotection, the ivDde-group was removed by reacting with a 5% hydrazine solution in DMF (v/v) for 5 minutes, twice. Then, lipoic acid was coupled to the lysine side chain (5eq. lipoic acid, 5eq. HATU, 10eq. DIPEA in DMF) for one hour, twice. After final Fmoc-deprotection, the peptide was cleaved from the resin in a mixture of TFA:TIPS:H_2_O:DMS at the ratio of 85:2.5:2.5:10 for 3 hours at room temperature. The peptide was precipitated using ice-cold diethyl ether, centrifuged at 4500 rpm at 4 °C for 5 minutes, re-dissolved in a mixture of MQ-H_2_O and MeCN and purified by semi-preparative reverse phase chromatography eluting with 5 to 95% MeCN in H_2_O over a C18 column. Peaks containing peptides were identified by LC-MS (MS calculated for C_40_H_69_N_9_O_14_S_2_ [M + 2H]^2+^, 428.7; observed 482.9), pooled, and concentrated in-vacuo to yield a white solid.

## Data Availability

All data supporting the findings of this study are available within the paper and its Supplementary Information. Source data for all figures are provided with this paper. The metabolomics and RNA-sequencing datasets generated in this study have been deposited in publicly accessible repositories, including the Gene Expression Omnibus (GEO; NCBI) and MetaboLights (EMBL-EBI); accession numbers will be provided upon acceptance. All other data are available from the corresponding author upon reasonable request.

## Supporting information

Ghosh et al. Extended Data

## Acknowledgements

We thank members of the Brady, Burslem, Cobine, DeNicola, Leary, Snyder, and Wellen, laboratories, the Penn Therapeutics Mechanisms Group, I. Asangani, W. Bailis,

T.P. Gade, B. Keith, M.C. Simon, D. Sneddon, P.M. Titchenell, S. Venkatesh, and K.E. Vest, for administrative, technical support, discussions, and/or review of the manuscript. This work was supported by NIH grants R35GM124749 (D.C.B.), R01CA280833 (D.C.B. & G.M.B), R35GM142505 (GMB), Pew Charitable Trusts Innovation Fund Award # 589950 (D.C.B.), Ludwig Institute for Cancer Research (D.C.B), R35GM156596 (N.W.S). Figures and graphical elements were created using BioRender.

## Contributions

S.G.: Conceptualization, Methodology, Investigation, Formal analysis, Data curation, Visualization. Performed Western blot analysis, Seahorse assays, ATP and NADH measurements, and molecular cloning AOX, NDI1, LbNOX, and mitoLbNOX, SLC25A3 WT and SLC25A3 mutant constructs, as well as COX, OGDH, PDH, and PC activity assays.

A.F.J.: Conceptualization, Methodology, Investigation, Formal analysis, Data curation, Visualization. Performed BAP probe labeling in lysates and co-immunoprecipitation experiments.

J.C.J.H.: Methodology, Investigation, Formal analysis. Synthesized BAP probe and Klip peptides and performed mass spectrometry analysis.

N.R.M.: Investigation. Performed CETSA experiments.

P.C.-P.: Investigation, Formal analysis. Performed ¹³C-glucose and ¹³C-glutamine tracing experiments and analyzed data.

N.H.N.: Investigation. Performed α-ketoglutarate and 2-hydroxyglutarate measurements.

Y.K.: Investigation, Formal analysis. Performed unbiased metabolomics and data analysis.

G.M.D.: Conceptualization, Supervision, Formal analysis. Contributed expertise in unbiased metabolomics, data analysis, and discussion.

A.J.: Investigation. Performed CoA measurements.

N.W.S.: Investigation, Formal analysis. Performed CoA measurements and contributed metabolomics expertise and analysis.

S.C.L.: Resources, Supervision. Provided *Slc25a3* MEFs and contributed expertise and discussion.

P.A.C.: Resources, Supervision. Provided *Slc25a3* mutants, MEFs, and contributed expertise and discussion.

C.R.B.: Investigation, Formal analysis. Performed α-ketoglutarate and 2-hydroxyglutarate measurements and contributed expertise and discussion.

K.E.W.: Conceptualization, Supervision, Formal analysis. Contributed expertise in isotope tracing, metabolic analysis, and discussion.

G.M.B.: Conceptualization, Methodology, Supervision, Formal analysis. Contributed to BAP probe and Klip synthesis, mass spectrometry strategy, and discussion.

D.C.B.: Conceptualization, Supervision, Funding acquisition, Writing – original draft, Writing – review & editing. Conceived the project, oversaw all aspects of the study, and contributed to study design and data analysis.

All authors: Writing – review & editing.

## Ethics declarations

No potential conflicts of interest were disclosed by the other authors.

---

**Extended Data Table 1.**
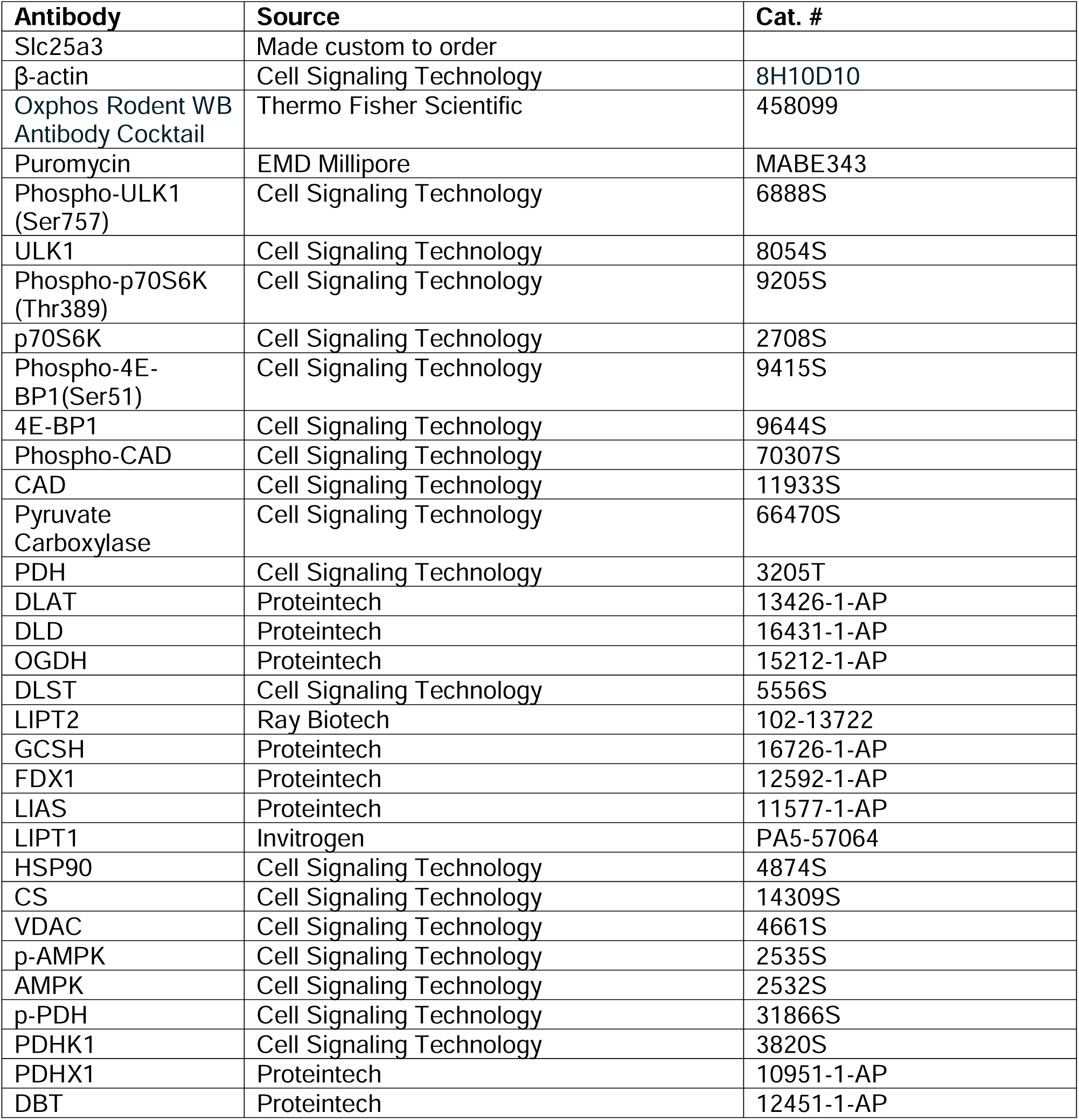

| **Antibody** | **Source** | **Cat. #** |
| --- | --- | --- |
| Slc25a3 | Made custom to order |  |
| β-actin | Cell Signaling Technology | 8H10D10 |
| Oxphos Rodent WB Antibody Cocktail | Thermo Fisher Scientific | 458099 |
| Puromycin | EMD Millipore | MABE343 |
| Phospho-ULK1 (Ser757) | Cell Signaling Technology | 6888S |
| ULK1 | Cell Signaling Technology | 8054S |
| Phospho-p70S6K (Thr389) | Cell Signaling Technology | 9205S |
| p70S6K | Cell Signaling Technology | 2708S |
| Phospho-4E-BP1(Ser51) | Cell Signaling Technology | 9415S |
| 4E-BP1 | Cell Signaling Technology | 9644S |
| Phospho-CAD | Cell Signaling Technology | 70307S |
| CAD | Cell Signaling Technology | 11933S |
| Pyruvate Carboxylase | Cell Signaling Technology | 66470S |
| PDH | Cell Signaling Technology | 3205T |
| DLAT | Proteintech | 13426-1-AP |
| DLD | Proteintech | 16431-1-AP |
| OGDH | Proteintech | 15212-1-AP |
| DLST | Cell Signaling Technology | 5556S |
| LIPT2 | Ray Biotech | 102-13722 |
| GCSH | Proteintech | 16726-1-AP |
| FDX1 | Proteintech | 12592-1-AP |
| LIAS | Proteintech | 11577-1-AP |
| LIPT1 | Invitrogen | PA5-57064 |
| HSP90 | Cell Signaling Technology | 4874S |
| CS | Cell Signaling Technology | 14309S |
| VDAC | Cell Signaling Technology | 4661S |
| p-AMPK | Cell Signaling Technology | 2535S |
| AMPK | Cell Signaling Technology | 2532S |
| p-PDH | Cell Signaling Technology | 31866S |
| PDHK1 | Cell Signaling Technology | 3820S |
| PDHX1 | Proteintech | 10951-1-AP |
| DBT | Proteintech | 12451-1-AP |

**Extended Data Figure 1.**
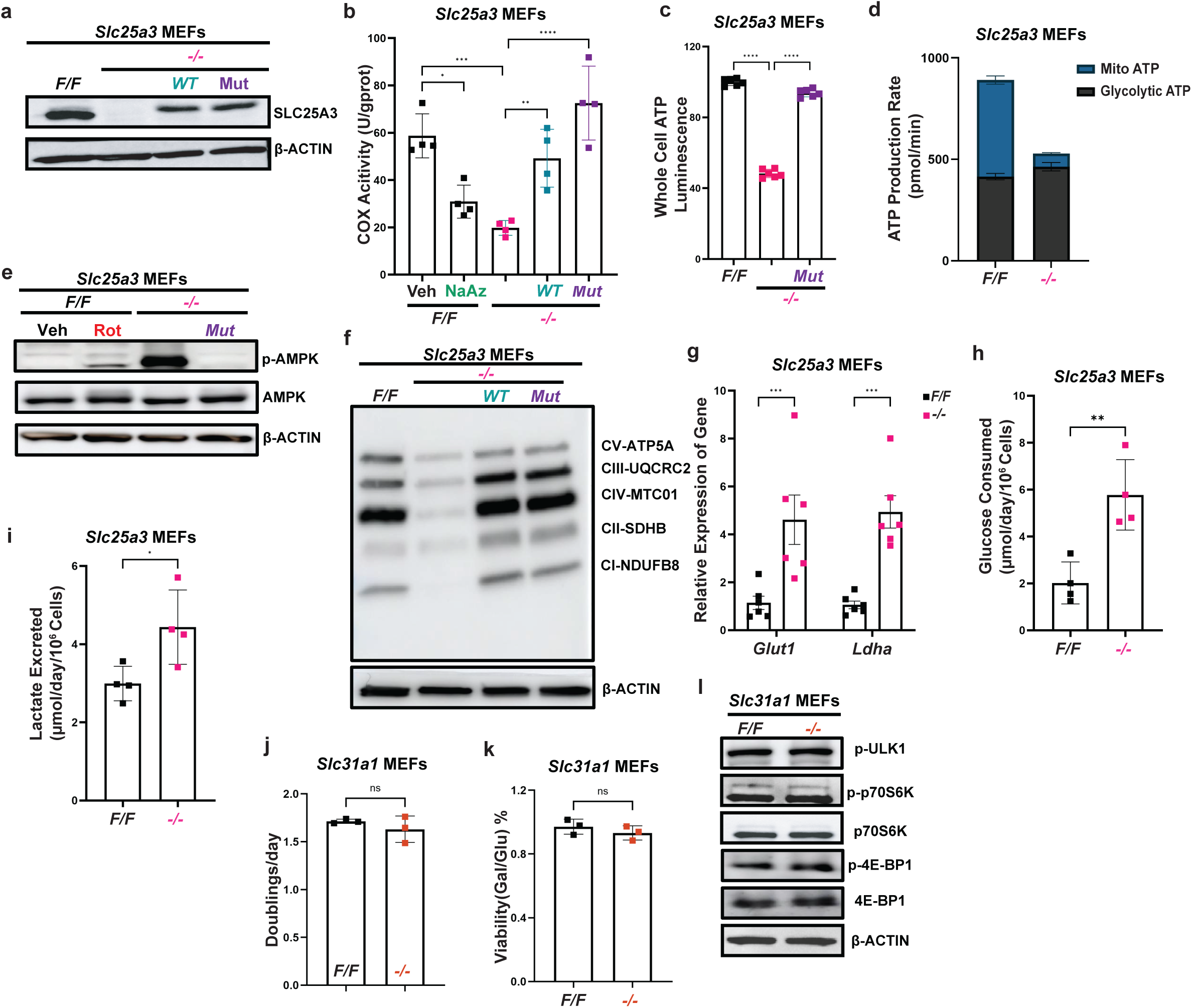
**a,** Immunoblot detection of *Slc25a3* or β-ACTIN in *Slc25a3 F/F*, *-/-*, human *WT Slc25a3* re-expressed into *-/-* (*WT*), and mutant *Slc25a3*(L175A) reinserted into *-/-*(*Mut*) mouse embryonic fibroblasts (MEFs). **b,** Cytochrome Oxidase activity measured using an activity assay kit in isolated mitochondria from *Slc25a3 F/F* MEFs treated with either vehicle or sodium azide (1µM), *Slc25a3 -/-*, *WT* and *Mut* MEFs, normalized to respective protein concentrations. **c,** Whole cell ATP levels measured in *Slc25a3 F/*F, *-/-* and *Mut* MEFs. **d,** ATP production rates by glycolytic and mitochondrial processes, with sequential injections of 1.5µM oligomycin and 0.5µM Rotenone/Antimycin A, measured with a Seahorse Analyzer, in *Slc25a3 F/F* and *-/-*MEFs. **e,** Immunoblot detection of p-AMPK, total AMPK or β-ACTIN in *Slc25a3 F/F* MEFs treated with either vehicle or 200nM Rotenone, *Slc25a3 -/-*, *Mut* MEFs. **f,** Immunoblot detection of the ETC components— ATP5A, MTC01, UQCRC2, SDHB, NDUFB8— or β-ACTIN in *Slc25a3 F/F*, *-/-*, *WT* and *Mut* MEFs. **g,** Relative Expression of mRNA levels of Glut1 and Ldha, represented as a fold change of *Slc25a3 -/-* vs *F/F* MEFs. **h, i,** Rate of **(h)** glucose consumed or **(i)** lactate excreted by *Slc25a3 F/F* or *-/-*MEFs, measured using a YSI metabolite analyzer. **j,** Doublings per day calculated from cell counts measured using trypan blue dye exclusion in Slc31a1 *F/F* or *-/-* MEFs. **k,** Viability (% of cells alive) of Slc31a1 *F/F* and *-/-* MEFs in 10mM Galactose or 10mM Glucose measured using trypan blue dye exclusion, represented as a ratio of viability in galactose over viability in glucose. **l,** Immunoblot detection of p-ULK1, p-p70S6K, total p70S6K, p-4E-BP1, total 4E-BP1, or β-ACTIN levels in Slc31a1 *F/F* or *-/-* MEFs. For **a-c**, **e-k**, n=3 biologically independent experiments, for **d** and **l**, n=1 biologically independent experiment. For **b-d**, **g-k**, data represented as mean ± SEM. ns= not significant, *p≤0.05, **p≤0.01, ***p≤0.001, ****p≤0.0001.

**Extended Data Figure 2.**
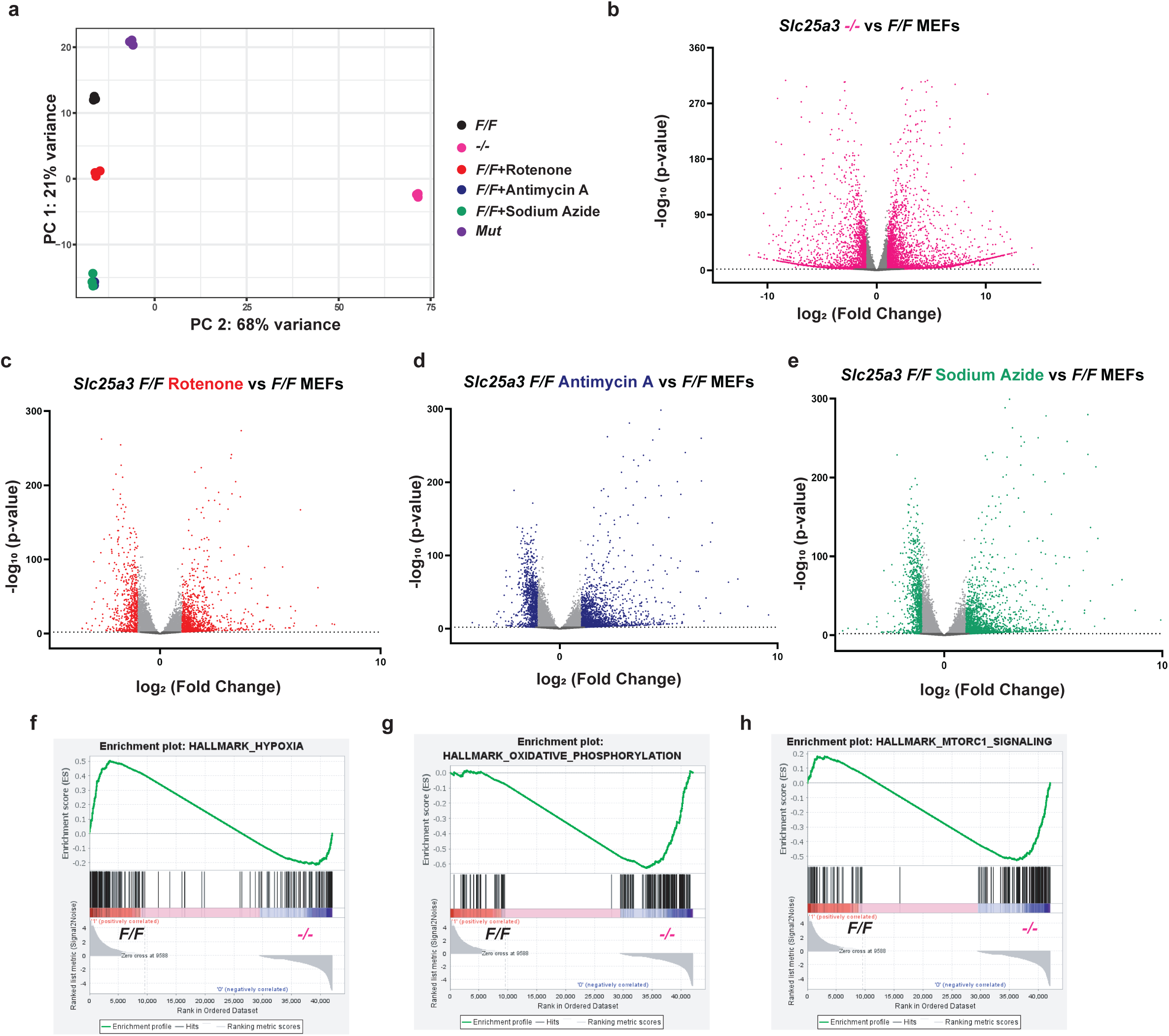
**a,** Principal component analysis of RNA-seq data from *Slc25a3 F/F* MEFs treated with vehicle (black circles), 200nM Rotenone for 24 hours (red circles), 10ng/ml Antimycin A for 24 hours (dark blue circles), 1µM Sodium azide for 24 hours (forest green circles), *-/-* MEFs (pink circles) or Mut MEFs (violet circles). PC1 (21% variance) separates treated from control samples, while PC2 (68% variance) captures residual donor variability. Count data were variance-stabilized transformed using DESeq2, and PCA was performed on the 500 most variable genes. **b, c, d, e,** Volcano plot summarizing differential expression analysis comparing **(b)** *Slc25a3 -/-*MEFs vs. *F/F* MEFs, **(c)** *Slc25a3 F/F* MEFs treated with 200nM rotenone for 24 hours vs. treated with vehicle, **(d)** *Slc25a3 F/F* MEFs treated with 10ng/ml Antimycin A for 24 hours vs. treated with vehicle, **(e)** *Slc25a3 F/F* MEFs treated with 1µM sodium azide for 24 hours vs. treated with vehicle. Genes with |log_2_ FC| > 1 and padj < 0.01 (DESeq2 Wald test) are considered significant (shown in respective colors). Upregulated genes (log_2_ FC > +1, padj < 0.01) are to the right; downregulated genes (log_2_ FC < -1, padj < 0.01) are to the left. Genes with padj < 0.01 but |log_2_ FC| ≤ 1 are shown in light gray — “statistically significant but not biologically significant”. Dark gray points represent non-significant genes (padj ≥ 0.01). The dashed horizontal line marks padj = 0.01 (-log_10_ = 2.0). All p-values were adjusted for multiple testing using the Benjamini-Hochberg method. **f, g, h,** Gene Set Enrichment Analysis running enrichment curves for the Hallmark **(f)** Hypoxia, **(g)** Oxidative Phosphorylation or **(h)** mTORC1 Signaling gene set in *Slc25a3 -/-* vs *F/F* MEFs. Each panel shows the ranked gene list (x-axis) and running enrichment score (y-axis). Vertical black ticks indicate gene positions. Positive ES peaks (curves that peak on the left indicate gene set enrichment in the experimental condition. Negative ES peaks (curves that bottom out on the right) indicate gene set enrichment in the control. NES values: **(f)**: NES = +.51; **(g)**: NES = -0.63; **(h)**: NES = - 0.52. Ranking metric: log_2_ fold change (DESeq2 Wald statistic).

**Extended Data Figure 3.**
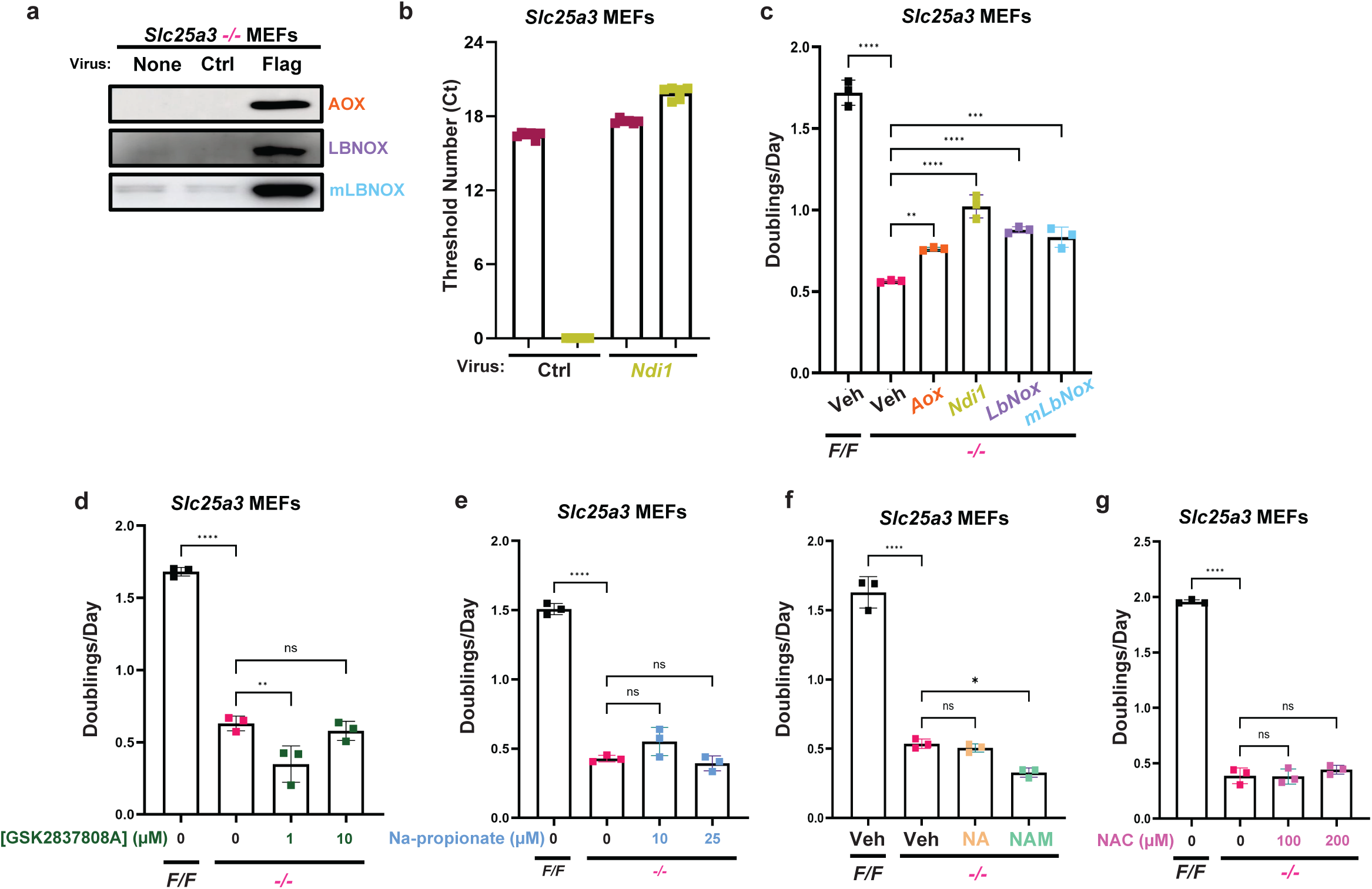
**a,** Immunoblot detection of flag-tagged AOX, LbNOX and mitoLbNox using flag antibody in *Slc25a3 -/-* MEFs transduced with no virus, a control virus corresponding to empty pCDH-puro vector, or virus produced from pCDH-puro-AOX-flag, pCDH-puro-LbNOX-flag or pCDH-puro-mitoLbNOX-flag viral plasmids. **b,** Cycle threshold values of Gapdh and Ndi1 in *Slc25a3 -/-* MEFs transduced with either a control virus corresponding to empty pCDH-puro vector or virus expressing pCDH-puro-NDI1. **c,** Doublings per day calculated from cell counts measured after 72 hours using trypan blue dye exclusion in *Slc25a3 F/F* MEFs treated with vehicle or *Slc25a3 -/-* MEFs treated with vehicle or transduced with AOX, NDI1, LbNox, or mitoLbNox virus. **d, e, f, g** Doublings per day calculated from cell counts measured using trypan blue dye exclusion in *Slc25a3 F/F* MEFs treated with vehicle or *Slc25a3 -/-* MEFs treated with either vehicle or **(d)** LDHa inhibitor, GSK2837808A (1µM or 10µM), **(e)** sodium propionate (10µM or 25µM), **(f)** nicotinic acid (10mM) or nicotinamide (1mM), **(g)** N-acetyl cysteine (100µM or 200µM) for 48 hours. For **c-g**, n=2 biologically independent experiments. For **b-g** data represented as mean ± SEM. ns= not significant, *p≤0.05, **p≤0.01, ***p≤0.001, ****p≤0.0001.

**Extended Data Figure 4.**
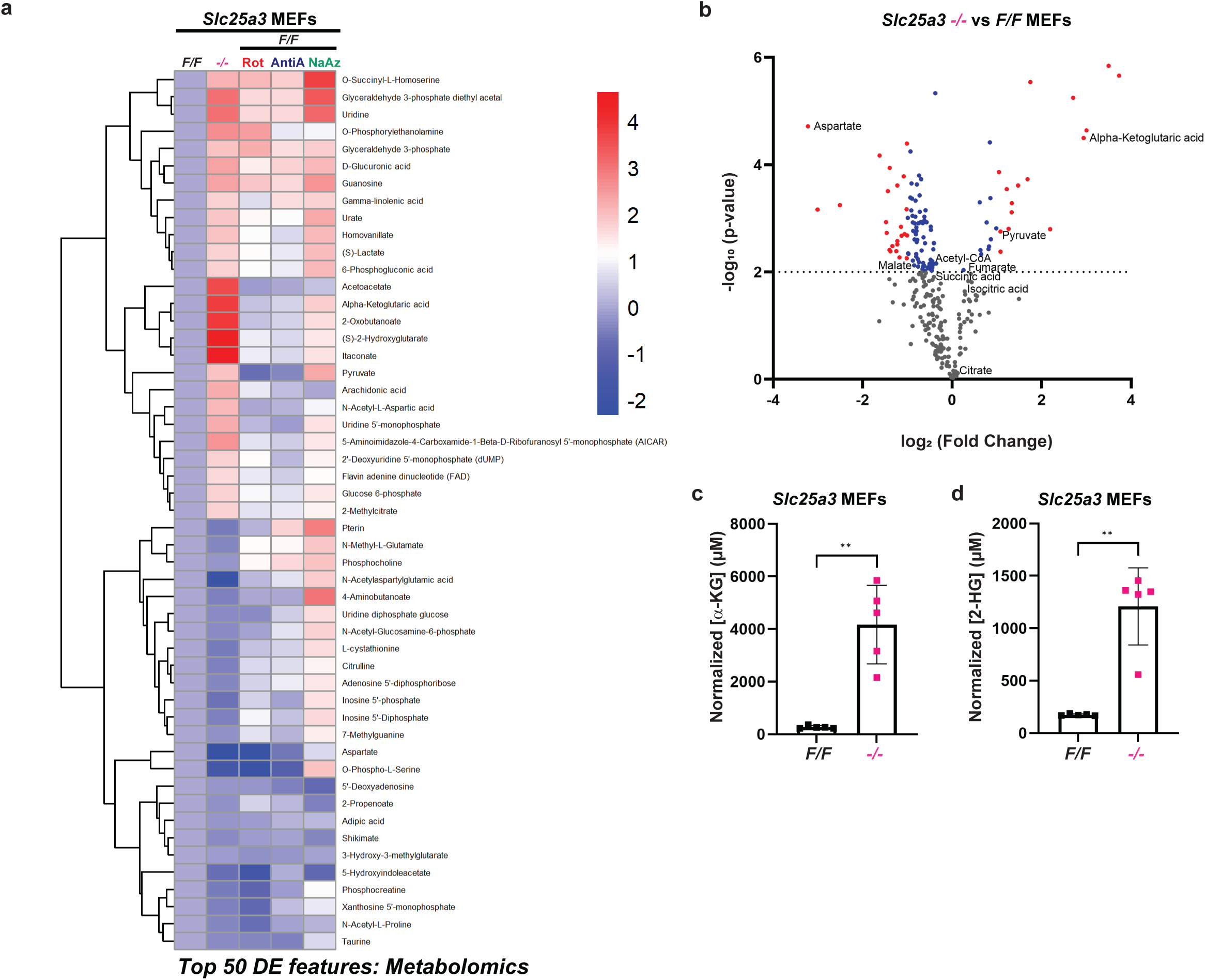
**a,** Heatmap of the top 50 differentially regulated metabolites in *Slc25a3 F/F* MEFs treated with 200nM Rotenone, 10ng/µl Antimycin A or 1µM Sodium Azide for 24 hours, or *Slc25a3 -/-* MEFs. The color bar represents z-scores. **b,** Volcano plot summarizing differential abundance analysis comparing *Slc25a3 -/-* vs *F/F* MEFs. Metabolites with |log_2_ FC| > 1 and padj < 0.01 are considered significant (shown in red). Upregulated metabolites (log_2_ FC > +1, padj < 0.01) are to the right; downregulated metabolites (log_2_ FC < -1, padj < 0.01) are to the left. Metabolites with padj < 0.01 but |log_2_ FC| ≤ 1 are shown in blue. Dark gray points represent non-significant changes in metabolites (padj ≥ 0.01). The dashed horizontal line marks padj = 0.01 (-log_10_ = 2.0). All p-values were adjusted for multiple testing using the Benjamini-Hochberg method. TCA cycle metabolites, acetyl CoA and aspartate are marked beside their respective points. **c, d** Concentration of **(c)** α-ketoglutarate (α-KG) and **(d)** 2-hydroxyglutarate (2-HG) in *Slc25a3 F/F* or *-/-* cells normalized to a standard, cell count and cell volume, measured using LC-MS. For **c and d,** n=1 biologically independent experiment, data represented as mean ± SEM. **p≤0.01.

**Extended Data Figure 5.**
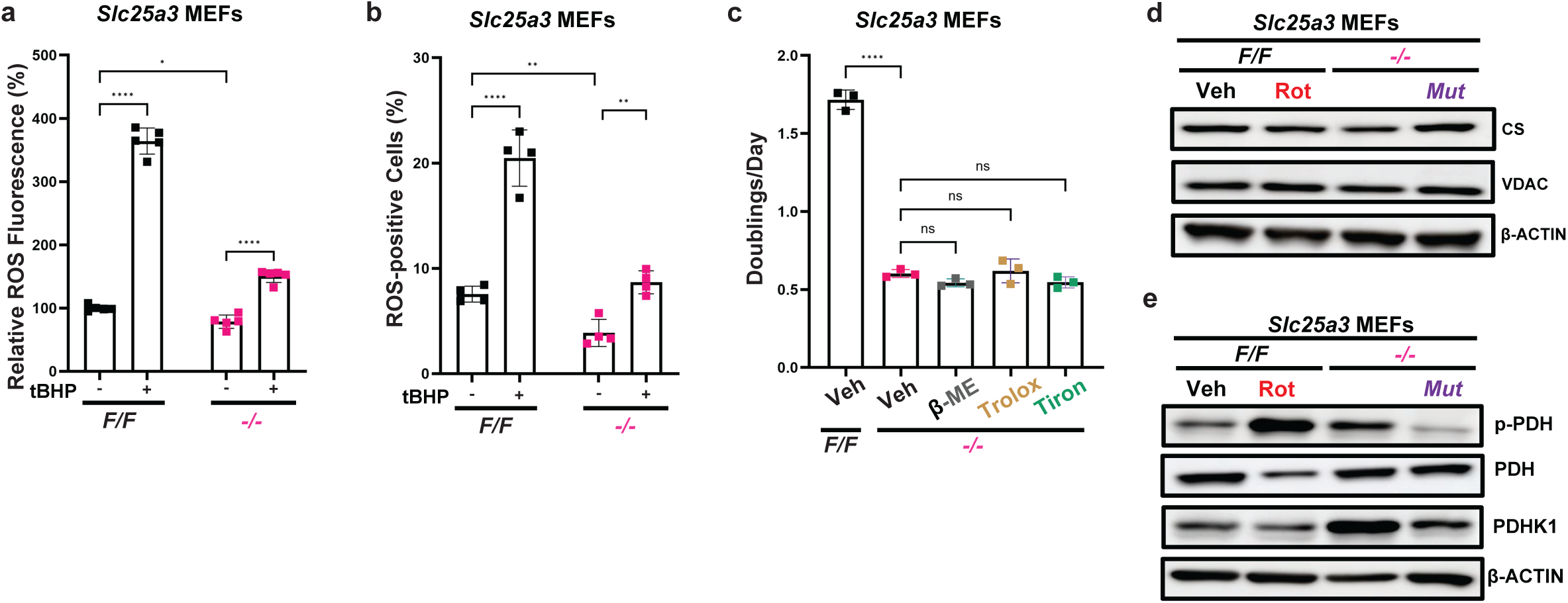
**a,** Relative reactive oxygen species (ROS) fluorescence levels measured with a ROS detection kit, in *Slc25a3 F/F* and *-/-* MEFs treated with either vehicle or 55µM tert-butyl hydroperoxide (tBHP) for 4 hours. **b,** Flow cytometric analysis of reactive oxygen species (ROS) levels in *Slc25a3 F/F* and *-/-* MEFs treated with either vehicle or 55µM tBHP for 4 hours. ROS were detected using the cell-permeable probe H_2_DCFDA (5 µM, 30 min at 37°C). ROS-positive cells were defined as the percentage of cells with fluorescence intensity exceeding a threshold set at the 99th percentile of unstained cells. **c,** Doublings per day calculated from cell counts measured using trypan blue dye exclusion in *Slc25a3 F/F* MEFs treated with vehicle or *Slc25a3 -/-*MEFs treated with either vehicle, 100µM β-mercaptoethanol (β-ME), 100µM Trolox or 250µM Tiron for 72 hours. **d, e,** Immunoblot detection in *Slc25a3 F/F* MEFs treated with either vehicle or 200nM Rotenone, *Slc25a3 -/-*, *Mut* MEFs of **(d)** Citrate synthase (CS), Voltage-dependent Anion Channel (VDAC) and β-ACTIN and **(e)** phospho-PDH (E1), total PDH (E1), PDH kinase1 (PDHK1) or β-ACTIN. For **a-e**, n=2 biologically independent experiments. For **a-c**, data represented as mean ± SEM. ns= not significant, *p≤0.05, **p≤0.01, ****p≤0.0001.

**Extended Data Figure 6.**
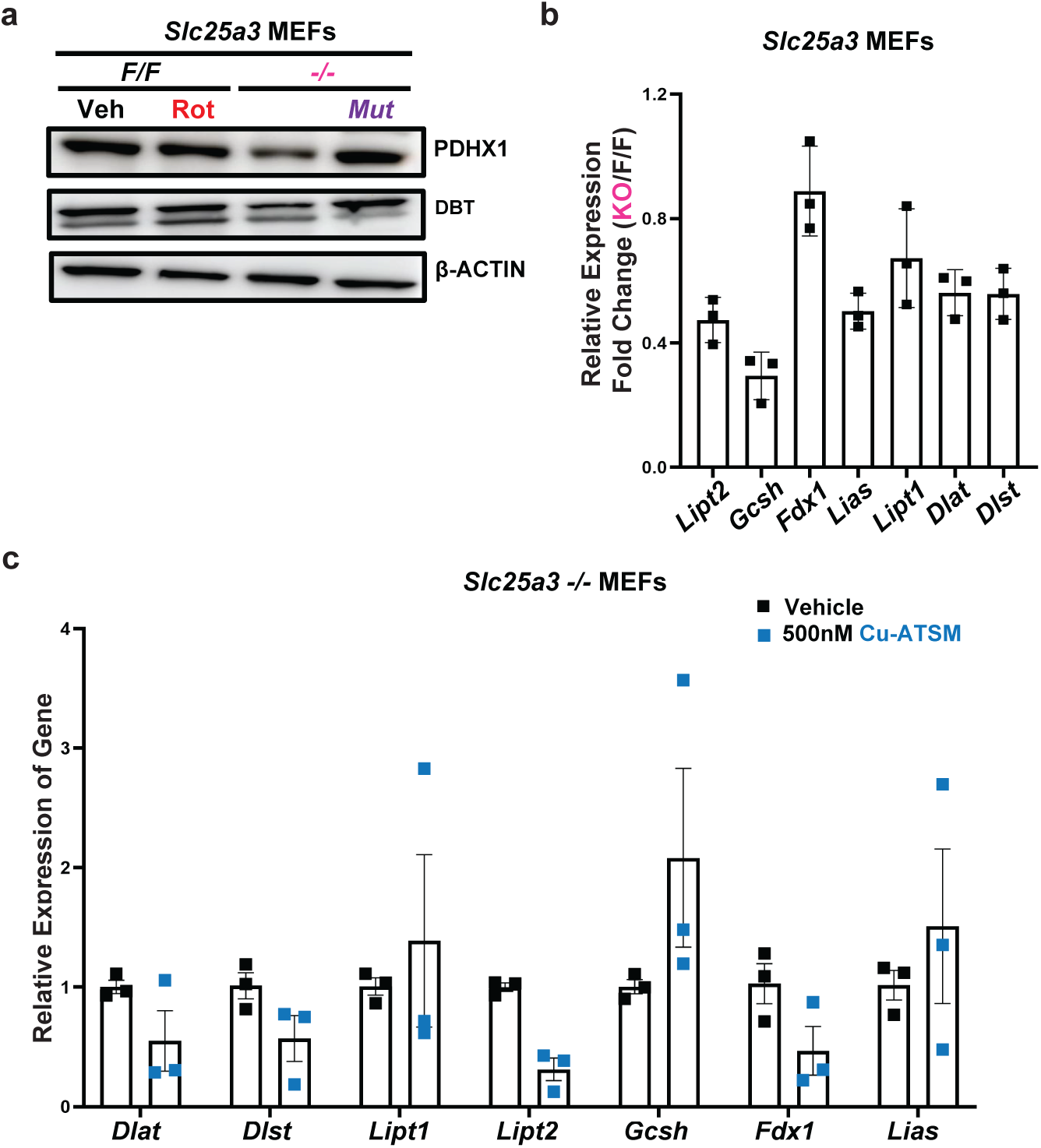
**a,** Immunoblot detection of Pyruvate Dehydrogenase Protein X Component, DBT or β-ACTIN in *Slc25a3 F/F* MEFs treated with either vehicle or 200nM Rotenone, *Slc25a3 -/-*, *Mut* MEFs. **b,** Relative Expression of mRNA levels of LIPT2, GCSH, FDX1, LIAS, LIPT1, DLAT and DLST, represented as a fold change of *Slc25a3 -/-* vs *F/F* MEFs. **c,** Normalized relative expression of mRNA levels of DLAT, DLST, LIPT1, LIPT2, GCSH, FDX1 and LIAS in *Slc25a3 -/-* MEFs treated with either vehicle or 500nM Cu-ATSM for 24 hours. For **a-c** n=1 biologically independent experiment. For **b** and **c**, data represented as mean ± SEM. ns= not significant.

**Extended Data Figure 7.**
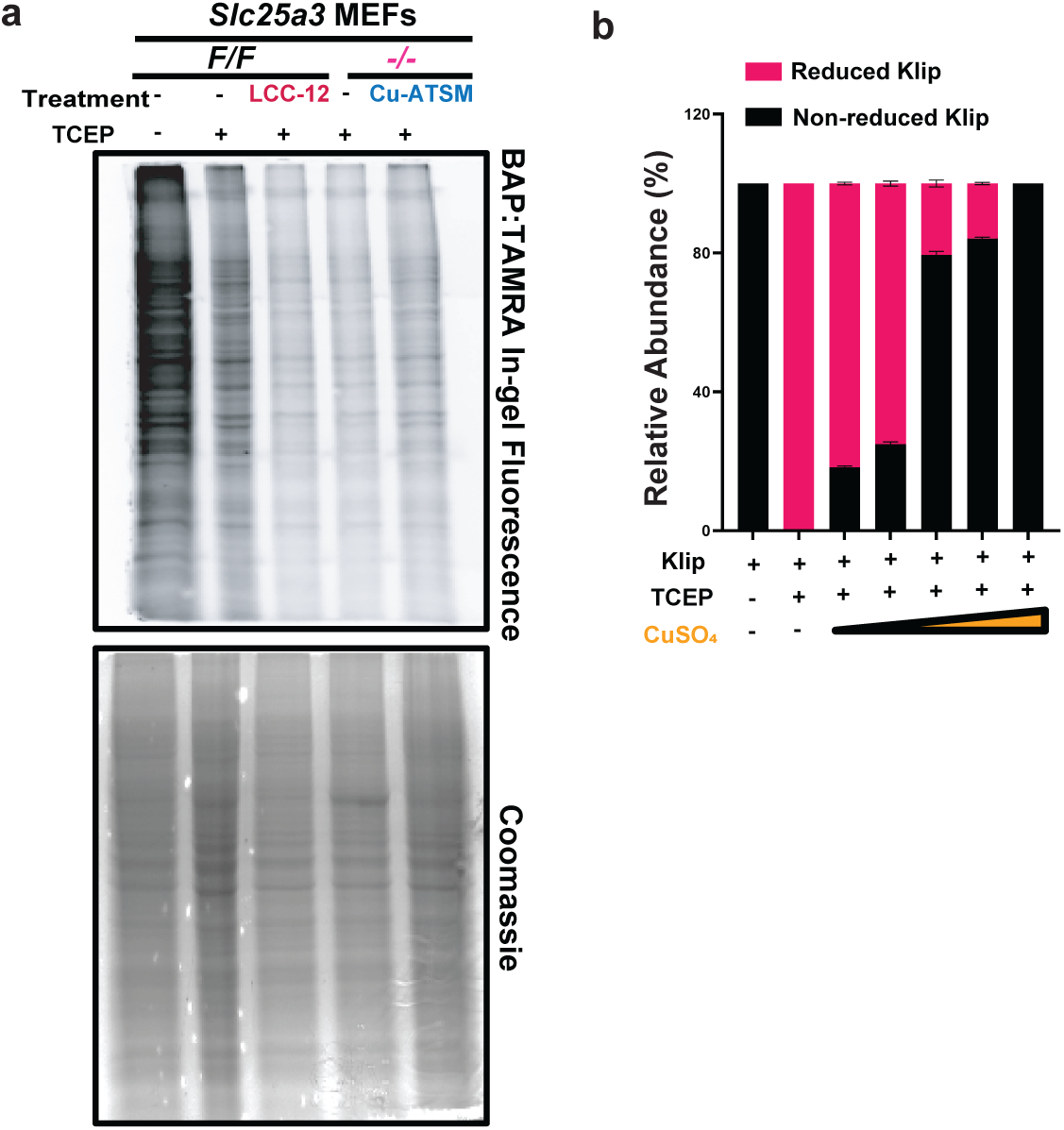
**a,** In-gel fluorescence imaging and Coomassie staining of *Slc25a3 F/F* MEFs treated with vehicle, lysates treated with TCEP (2mM) for 1 hour, or LCC-12 (30µM) for 24 hours and those lysates treated with TCEP (2mM) for 1 hour, and *Slc25a3 -/-* MEFs treated with either vehicle or Cu-ATSM (250nM) for 24 hours and those lysates treated with TCEP (2mM) for 1 hour. **b,** Relative abundance of the following species: DLAT(246-252)K249lip, retention time ∼7.05 min, [M+H]+ 964.44; DLAT(246-252)K249lip reduced, retention time ∼7.14 min, [M+H]+ 966.46 and DLAT(246-252)K249lip BAP conjugate, retention time ∼8.14 min, [M+H]+ 1044.50 detected using MS in reactions containing Klip only, Klip reduced with TCEP and increasing concentrations of CuSO_4_ (2.5, 5, 10, 25µM) for 4 hours. For **a,** n=3 biologically independent experiments. For **b,** n=3 independent replicates. For **b,** data represented as mean ± SEM.

