## Supplementary material for "Mitochondrial copper stabilizes lipoylated TCA cycle proteins to sustain metabolism and proliferation": Ghosh et al. Extended Data

Ghosh et al. Extended Data Figure 1

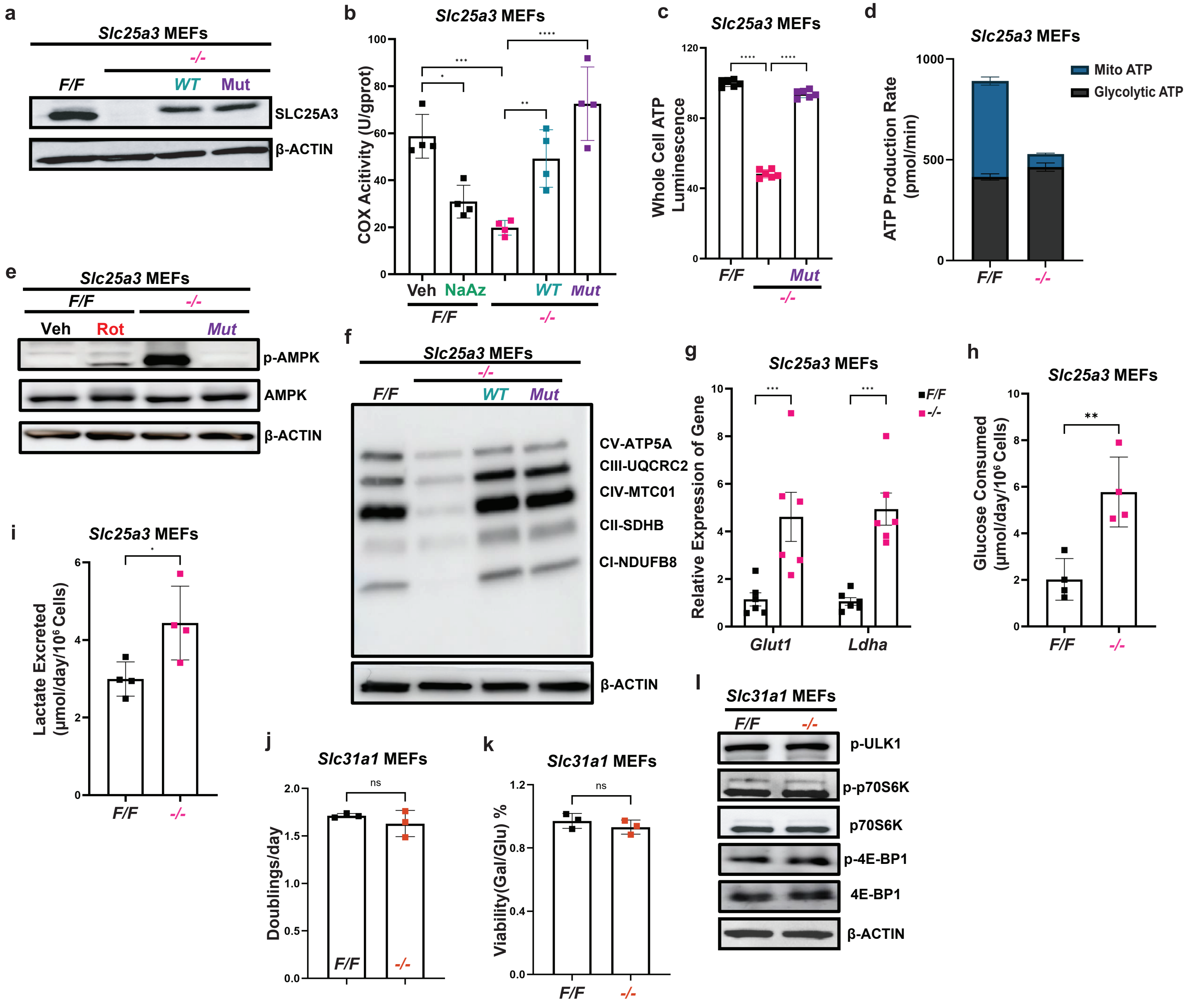

*Ghosh et al. Extended Data Figure 2*

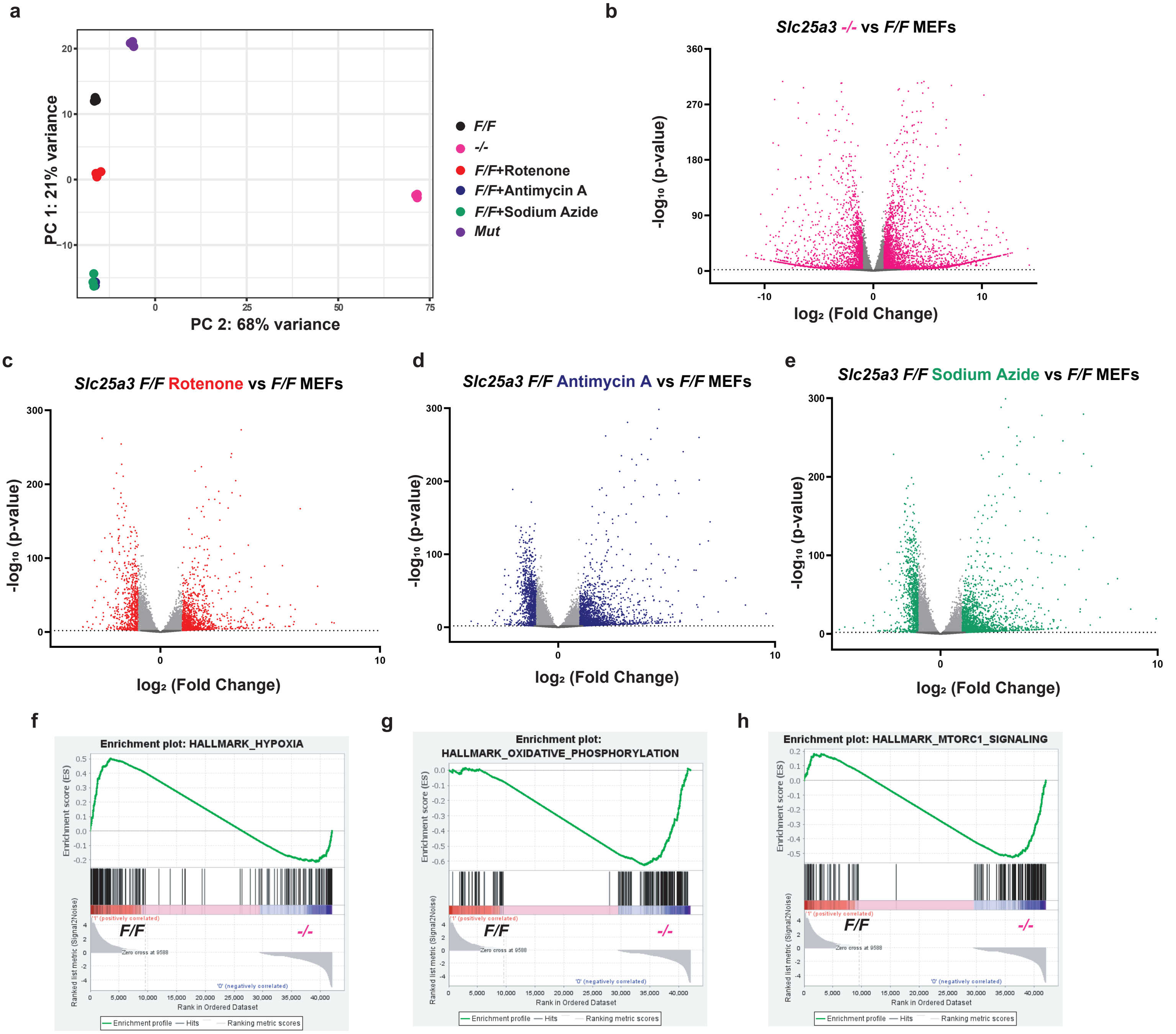

Ghosh et al. Extended Data Figure 3

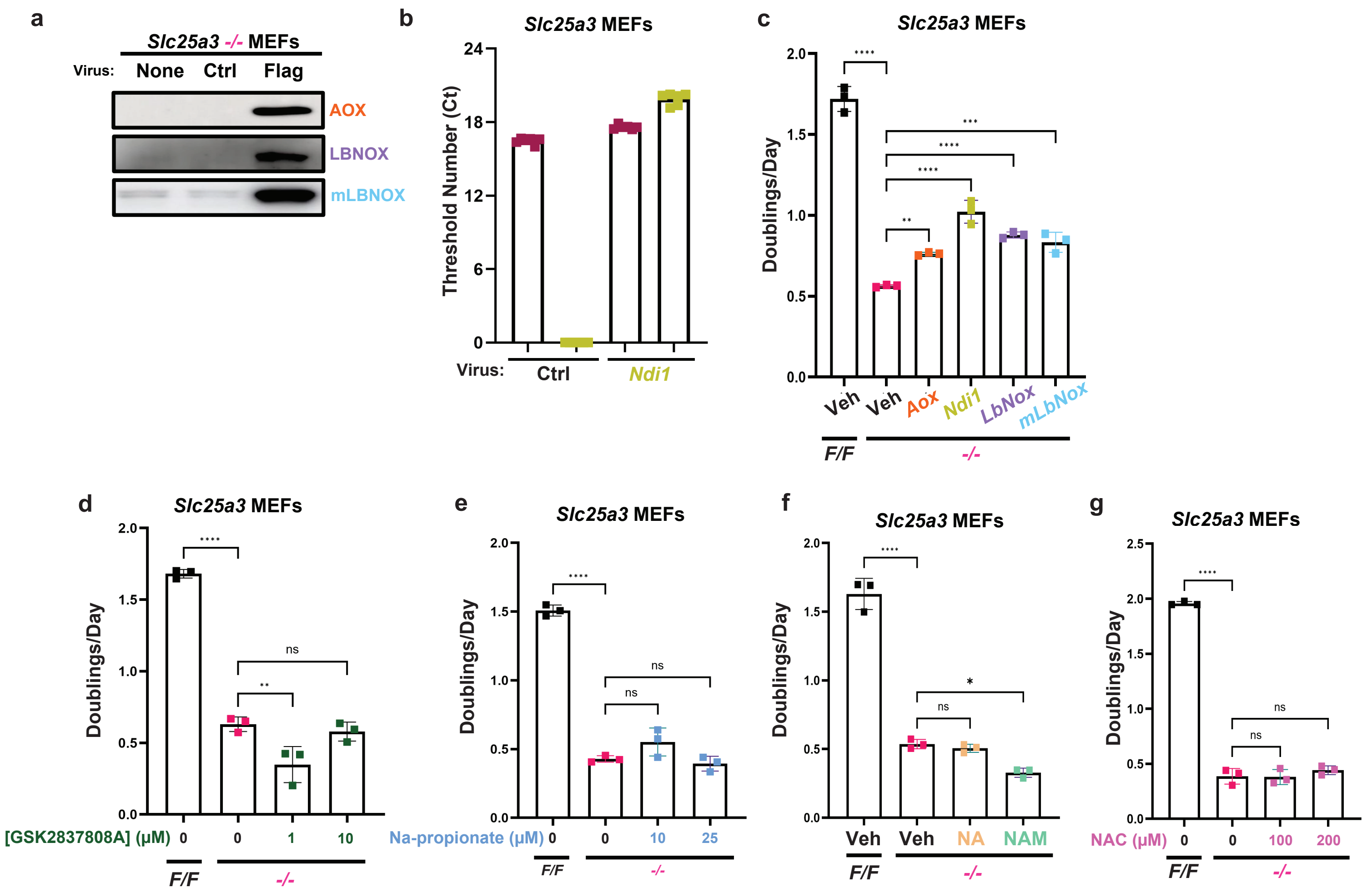

*Ghosh et al. Extended Data Figure 4*

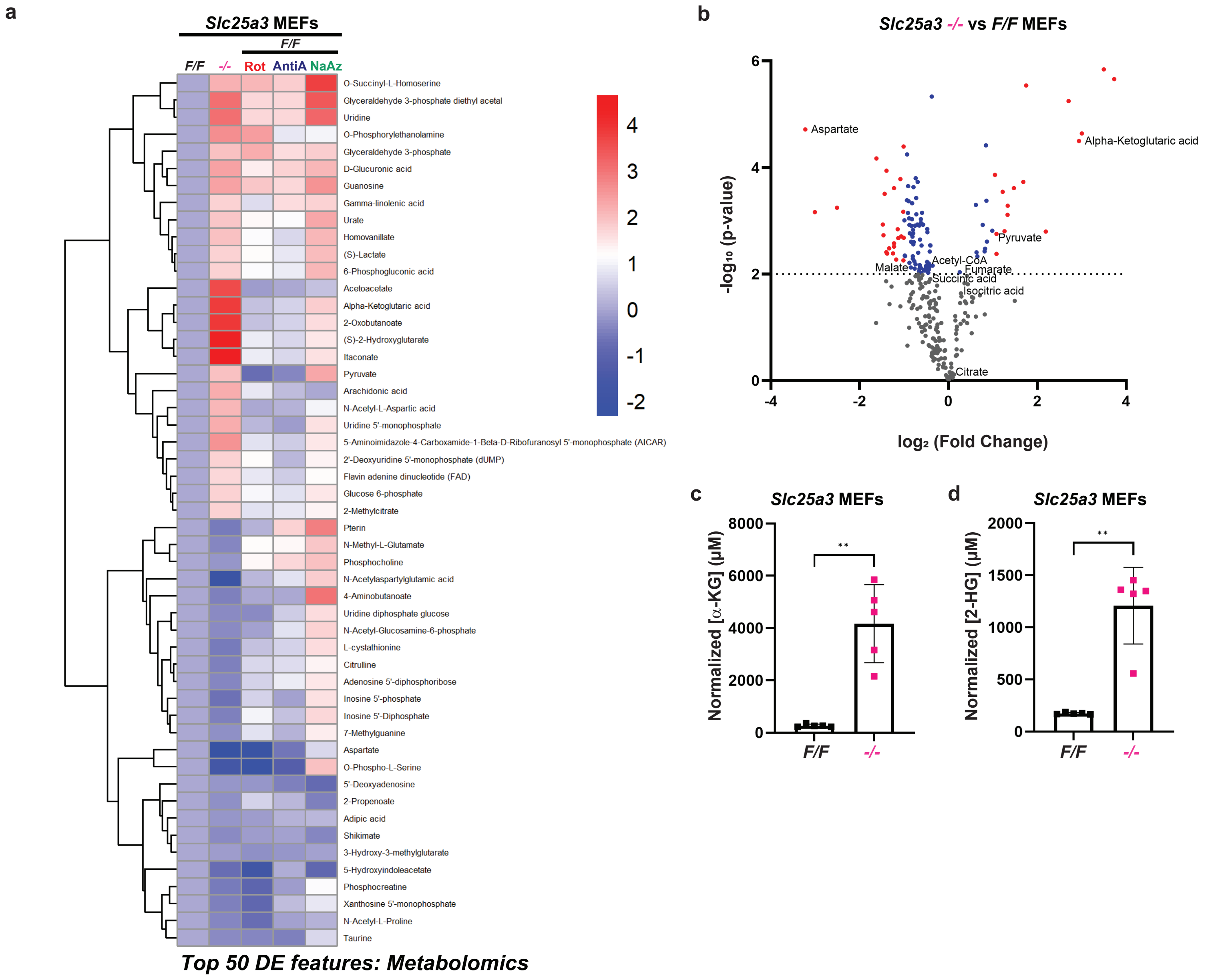

### Ghosh et al. Extended Data Figure 5

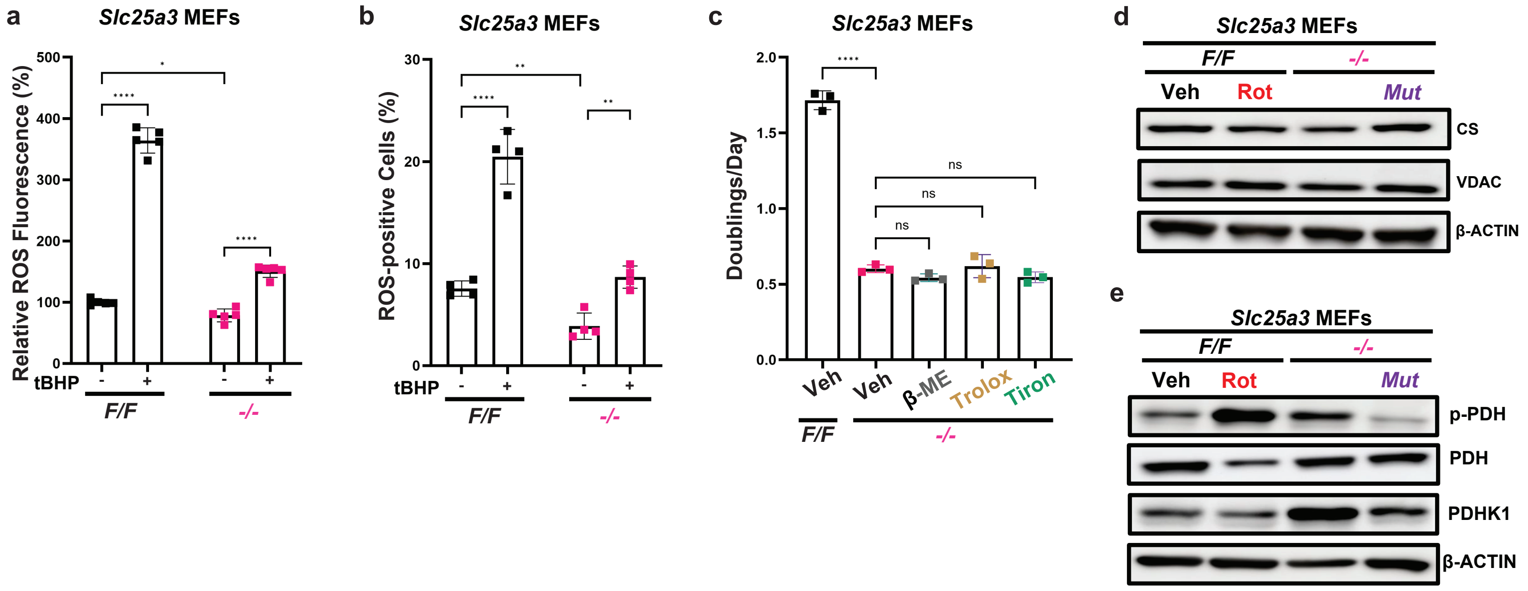

*Ghosh et al. Extended Data Figure 6*

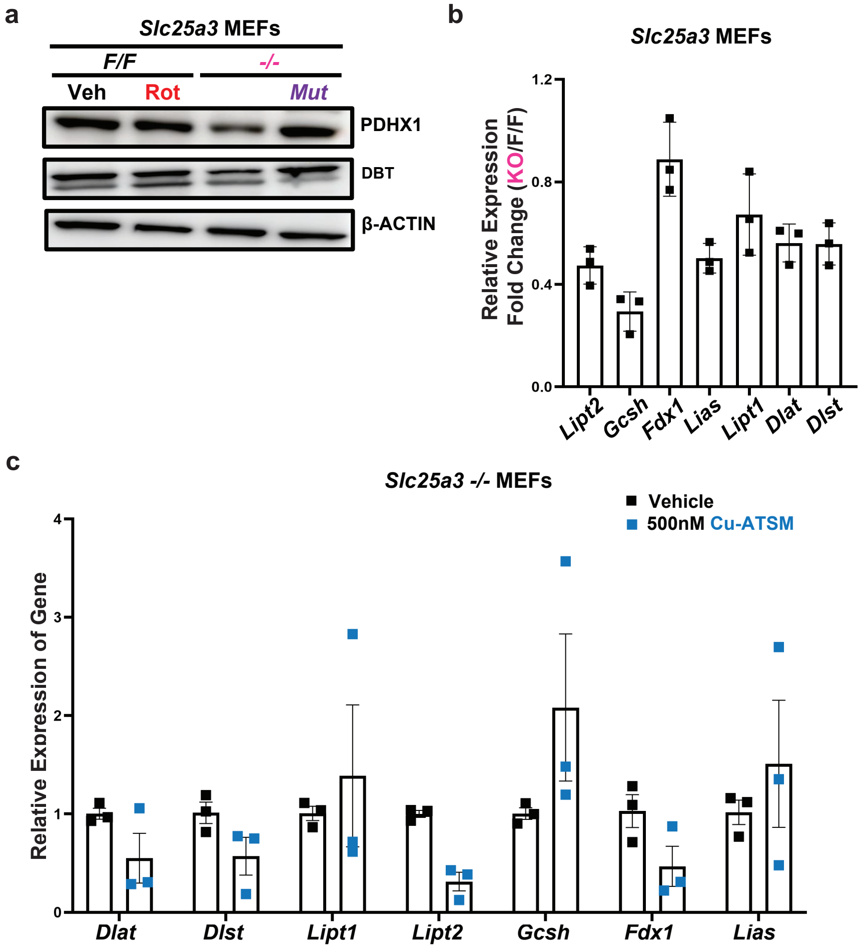

*Ghosh et al. Extended Data Figure 7*

a

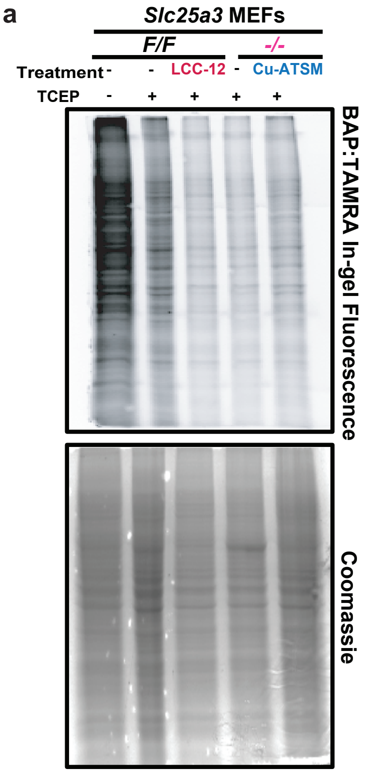


b

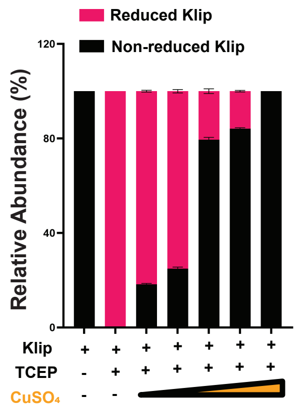
